# Identification of a divalent metal transporter required for cellular iron metabolism in malaria parasites

**DOI:** 10.1101/2024.05.10.587216

**Authors:** Kade M. Loveridge, Paul A. Sigala

## Abstract

*Plasmodium falciparum* malaria parasites invade and multiply inside red blood cells (RBCs), the most iron-rich compartment in humans. Like all cells, *P. falciparum* requires nutritional iron to support essential metabolic pathways, but the critical mechanisms of iron acquisition and trafficking during RBC infection have remained obscure. Parasites internalize and liberate massive amounts of heme during large-scale digestion of RBC hemoglobin within an acidic food vacuole (FV) but lack a heme oxygenase to release porphyrin-bound iron. Although most FV heme is sequestered into inert hemozoin crystals, prior studies indicate that trace heme escapes biomineralization and is susceptible to non-enzymatic degradation within the oxidizing FV environment to release labile iron. Parasites retain a homolog of divalent metal transporter 1 (DMT1), a known mammalian iron transporter, but its role in *P. falciparum* iron acquisition has not been tested. Our phylogenetic studies indicate that *P. falciparum* DMT1 (PfDMT1) retains conserved molecular features critical for metal transport. We localized this protein to the FV membrane and defined its orientation in an export-competent topology. Conditional knockdown of PfDMT1 expression is lethal to parasites, which display broad cellular defects in iron-dependent functions, including impaired apicoplast biogenesis and mitochondrial polarization. Parasites are selectively rescued from partial PfDMT1 knockdown by supplementation with exogenous iron, but not other metals. These results support a cellular paradigm whereby PfDMT1 is the molecular gatekeeper to essential iron acquisition by blood-stage malaria parasites and suggest that therapeutic targeting of PfDMT1 may be a potent antimalarial strategy.

## INTRODUCTION

Malaria is one of the deadliest infectious diseases worldwide and kills over 600,000 people yearly, with the highest mortality in children under five (1). The most virulent form of malaria is caused by *Plasmodium falciparum* parasites, which are responsible for most malaria deaths. The clinical manifestations of malaria occur when parasites invade red blood cells (RBCs), causing blood flow obstruction, endothelial damage, and inflammation that can lead to life-threatening end-organ failure (2). Rising parasite resistance to front-line artemisinin combination therapies poses a significant threat to global malaria control efforts (3). Understanding the essential parasite mechanisms for growth and replication within RBCs is an urgent challenge, and uncovering these survival strategies is key to developing innovative treatments to combat malaria.

*P. falciparum*, like all cellular organisms, requires labile iron to synthesize protein cofactors such as iron-sulfur clusters (Fe-S) and di-iron centers that are essential for core metabolic processes such as mitochondrial respiration and DNA synthesis/repair (4). Malaria parasites must scavenge iron from the human host to satisfy their nutritional requirements within the infected RBC, but mechanisms for iron acquisition are largely undefined. Nevertheless, it has been known for decades that blood-stage malaria parasites are highly sensitive to iron chelators such as deferoxamine (DFO, EC_50_ values of ∼10 µM) (5). Multiple studies have provided evidence that iron chelators must be internalized inside parasites to exert their anti-*Plasmodium* activity. These prior observations strongly suggest that parasites rely on a labile iron pool generated inside the parasite rather than utilizing exchangeable iron in the serum and RBC cytosol (6–9). Understanding the source and location of this labile iron pool and the mechanisms blood-stage parasites use to access it remains a critical and unmet challenge.

The RBC contains abundant non-exchangeable iron in the form of hemoglobin (Hb). As parasites grow inside RBCs, copious amounts of Hb are internalized within an acidic food vacuole (FV) and degraded to release free heme. *P. falciparum* lacks an active heme oxygenase to cleave the porphyrin macrocycle and release iron (10, 11), and most heme liberated in the FV is biomineralized into chemically inert hemozoin crystals to neutralize cellular toxicity (12). Prior studies provide indirect evidence that a small portion of heme escapes biomineralization in the FV and can be mobilized by parasites to satisfy essential heme requirements. Indeed, *Plasmodium* requires heme for cytochrome function in the mitochondrial electron transport chain (13), but de novo heme synthesis is dispensable in blood-stage parasites (14–16). These observations indicate that parasites can scavenge host heme by an undefined mechanism to meet metabolic needs. Within the oxidizing FV environment, a trace amount of free heme is expected to be non-enzymatically degraded through peroxide-coupled oxidation that non-specifically cleaves the porphyrin ring and release a small pool of labile iron (17–19). The RBC also contains other iron metalloproteins (e.g., remnant ferritin, catalase) that are likely internalized into the FV and may serve as alternative sources of iron in this acidic organelle (20–22). Based on these observations, we considered the FV to be a likely source of labile iron for *P. falciparum* during RBC infection. This function for the parasite FV would be consistent with the role of similar acidic organelles found in yeast and mammals (e.g., the vacuole and lysosome), which act as labile iron storage compartments (23, 24).

Labile iron generated in the parasite FV would be expected to depend on a protein transporter for export and utilization in metabolic processes outside the FV. Homologs of the ZIPCO (Zrt-, Irt-like Protein domain-containing protein) (25) and VIT (vacuolar iron transporter) (26) iron transporters have been previously studied in *P. falciparum*, but each is dispensable for blood-stage parasite growth and has localization and/or gene expression profiles that are incompatible with a dominant role in iron acquisition by blood-stage parasites. *Plasmodium* also encodes a homolog of the high-affinity iron transporter, divalent metal transporter 1 (DMT1, Pf3D7_0523800), which was recently localized to the FV membrane (27, 28). DMT1 homologs are ubiquitous divalent transition metal transporters that are conserved across all domains of life (29). These transporters predominantly localize to acidic environments such as the mammalian lysosome or yeast vacuole, where they harness proton gradients to drive iron transport into the cytosol (30, 31). Multiple studies have observed that *P. falciparum* DMT1 (PfDMT1) is refractory to disruption, which provides indirect evidence for an essential function (27, 32). Studies in rodent-infecting *Plasmodium yoelii* have reported that moderate decreases in PyDMT1 transcripts reduced parasite growth but cannot distinguish if DMT1 is rigorously essential and/or partially redundant with other mechanisms of iron acquisition in blood-stage *Plasmodium* parasites (28).

We introduced a GFP-tagged PfDMT1 episome and edited the native PfDMT1 locus via CRISPR/Cas9 for localization and knockdown studies. Sequence, phylogenetic, and molecular topology analyses confirmed that PfDMT1 has conserved biochemical features for metal transport and a membrane orientation similar to other organisms. Conditional PfDMT1 knockdown (KD) caused rapid parasite death and disrupted iron-dependent processes, including apicoplast biogenesis and the mitochondrial electron transport chain. Iron supplementation rescued parasite growth from partial loss of PfDMT1 but not stringent KD, underscoring a dominant role for PfDMT1 in iron uptake by blood-stage malaria parasites. These findings reveal an essential molecular mechanism of iron scavenging in *P. falciparum* and suggest that PfDMT1 is a promising candidate for antimalarial drug development.

## RESULTS

### *Plasmodium* encodes a DMT1 homolog with conserved sequence and structural features

Multiple prior studies have noted that Pf3D7_0523800 shows general sequence similarity to human DMT1 (Uniprot P49281) (12, 27, 28). Using the human protein as query sequence, we conducted a BLAST search *of P. falciparum* and confirmed that Pf3D7_0523800 is the only DMT1 homolog detected in the parasite genome. To determine if PfDMT1 retained the expected sequence features of a metal transporter, we aligned PfDMT1 with other well-studied DMT1 transporters. PfDMT1 shares 29% sequence identity with human DMT1 and 22% identity with the yeast homolog (Uniprot Q12078), Smf3 (Fig. S1). PfDMT1 retains the essential DPGN motif on the first transmembrane domain (TMD), which is required for metal binding and transport in other organisms and involves direct coordination of metal ions by the Asp and Asn residues (Fig 1A.)(33). Curiously, PfDMT1 substitutes two conserved metal-binding residues on TMD 6— Ala with a Ser (carbonyl backbone coordination) at position 430 and a Met with a Ser at position 433. Analysis of prokaryotic DMT1 homologs has provided evidence that the Met residue on TMD 6 acts as a soft base that enables DMT1 transporters to distinguish softer transition metal ions like Fe^2+^ from hard alkaline metal ions like Ca^2+^. Mutations of the metal-coordinating Met ablated metal transport in some organisms, but in other cases, it permitted calcium as a substrate without impacting transition metal transport(34). The functional consequences of this Met-Ser replacement in PfDMT1 and its impact on transport specificity and other transport properties are uncertain.

**Figure 1.**
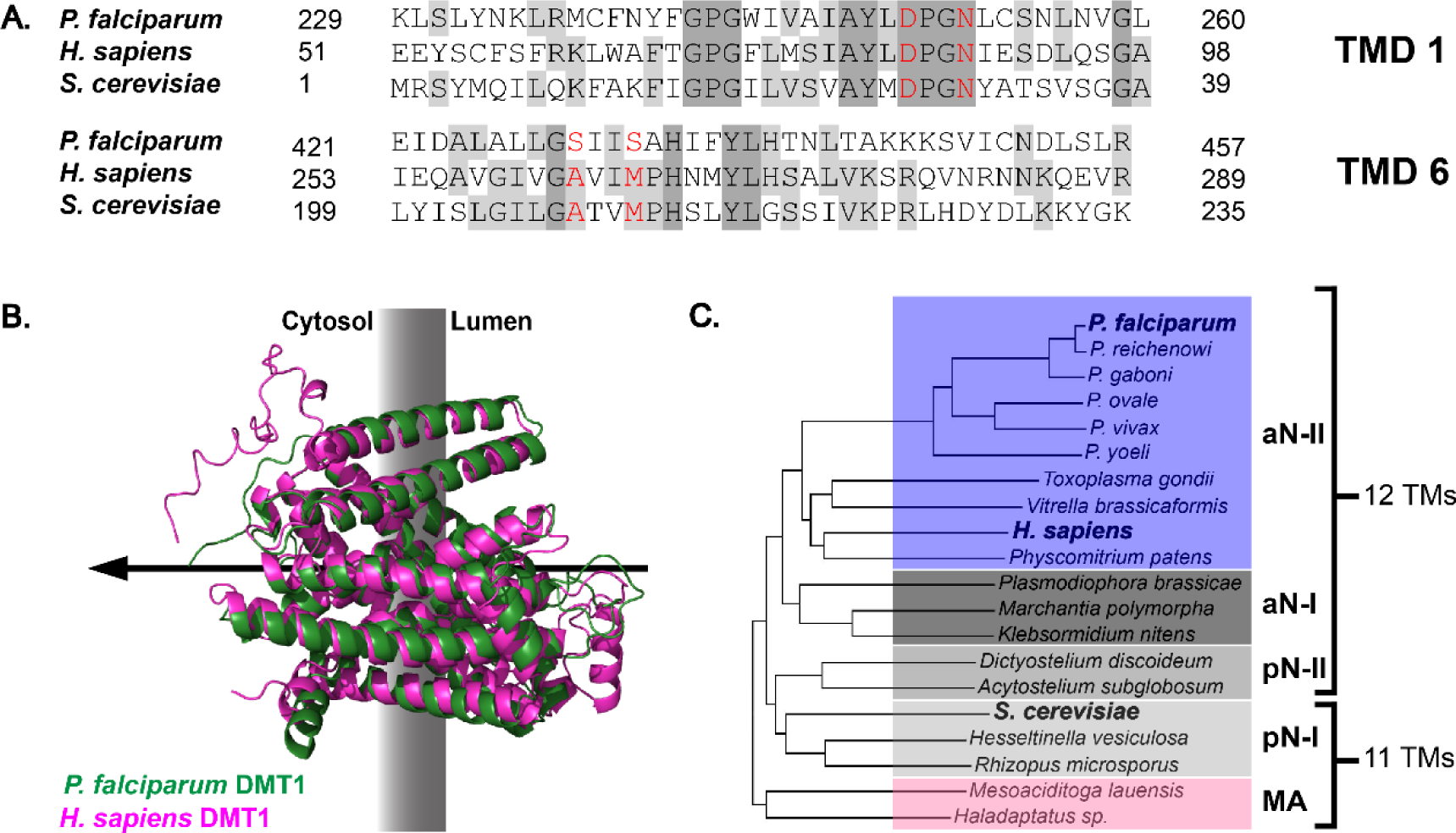
Sequence and structural properties of DMT1 homologs. **(A)** Sequence alignment of DMT1 homologs from *P. falciparum*, *H. sapiens*, and *S. cerevisiae* focusing on transmembrane domains 1 and 6 that are primarily responsible for metal coordination. The red amino acids are predicted to function as metal-binding ligands based on solved crystal structures of bacterial homologs. Note that the backbone carbonyl groups of *P. falciparum* S430, *H. sapiens* A262, and *S. cerevisiae* A208 are predicted to coordinate metals. Full sequence alignment is provided in Fig. S1. **(B)** Superposition of predicted AlphaFold structures of *H. sapiens* DMT1 and PfDMT1. The disordered N-terminus of each protein was excluded. The black arrow indicates the predicted direction of metal transport into the cytosol based on known properties of mammalian and yeast DMT1. **(C)** Simplified phylogenetic tree of *Plasmodium* homologs relative to other DMT1 transporters. DMT1 homologs are classified as archetype-I/II or prototype-I/II based on prior classifications or through grouping with the respective evolutionary subgroup. Prokaryotic MntH clade A homologs were used as the outgroup. A more detailed analysis is provided in Fig. S2.

No experimental structures have been determined for a eukaryotic DMT1 homolog. However, alignment of the AlphaFold-predicted PfDMT1 model with X-ray crystal structures of prokaryotic homologs revealed high structural similarity (Fig. S2) (35–37). A comparison of the AlphaFold structures of PfDMT1 and human DMT1 also suggested similar protein architectures (Fig. 1B). The PfDMT1 AlphaFold structure predicts 12 TMDs, which align with the 12 TMDs observed in the *Eremococcus colecola* DMT/MntH crystal structure (37) and in the human DMT1 AlphaFold structure (Fig. 1B and Fig. S2C). We noted that the Phobius transmembrane identifier tool also predicted that PfDMT1 contains 12 TMDs (38).

Previous phylogenetic analyses demonstrated that PfDMT1 shares ancestry with metazoan DMT1 transporters (39). We extended these prior analyses of PfDMT1’s subtype utilizing more detailed evolutionary classifications of DMT1 transporters that have emerged in recent years (40). We determined that PfDMT1 co-segregates with mammalian DMT1 homologs in the archetype-II group (Fig. 1D and Fig. S3) (40) that contain 12 TMDs (Fig. 2A), consistent with AlphaFold and Phobius predictions. Within this group, PfDMT1 exhibits higher sequence divergence from other archetype-II DMT1 homologs than closely related *Toxoplasma gondii* parasites (Fig. S3). PfDMT1 segregation with archetype-II homologs contrasts with yeast Smf3, which segregates with a more distant group of DMT1 homologs that are distinguished by 11 TMDs. The predicted 12 TMDs and conserved metal-binding motifs in PfDMT1 suggest a key role in metal transport at the parasite FV membrane analogous to DMT1 homologs in mammals and yeast.

**Figure 2.**
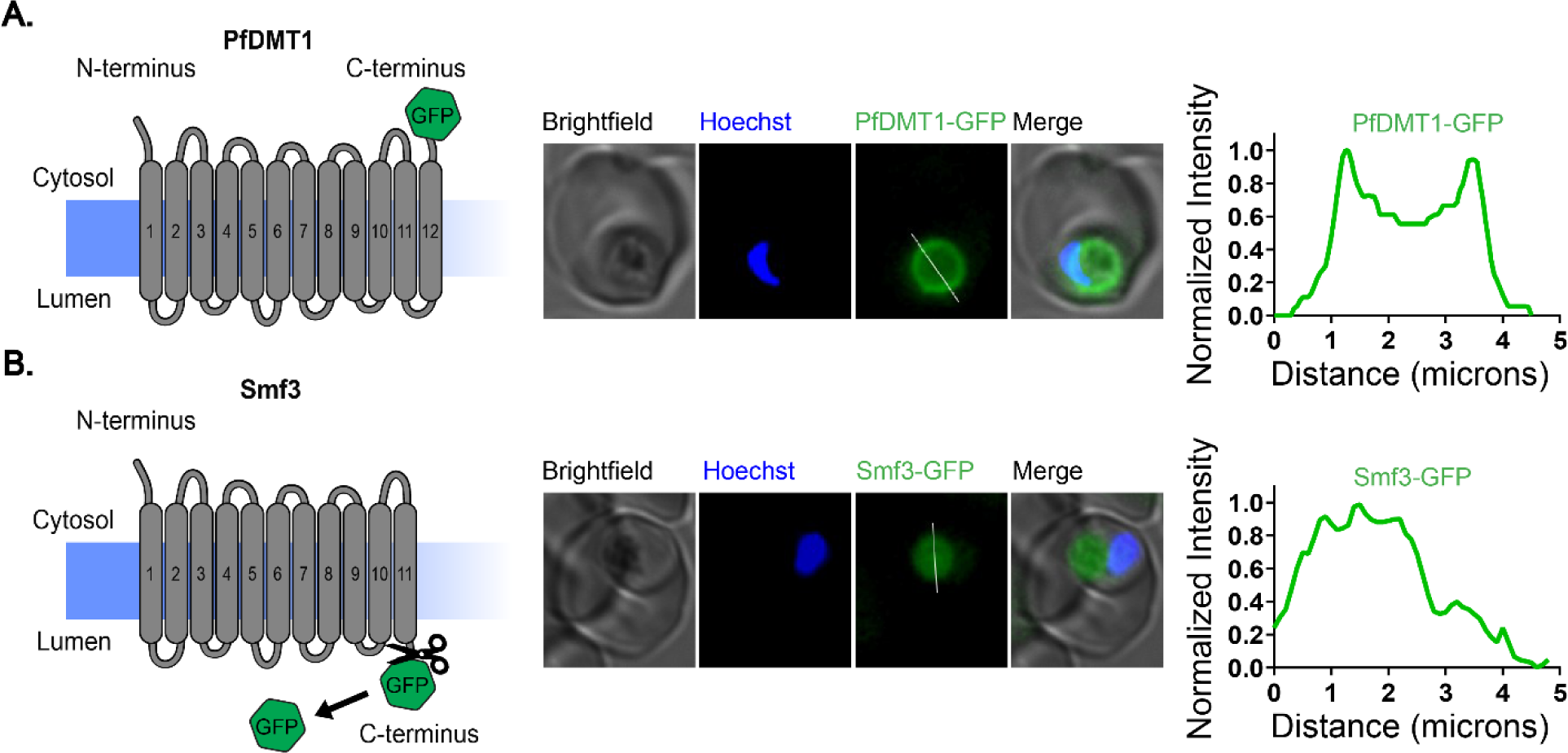
Orientation of DMT1 homologs in the FV membrane. Live-parasite fluorescence microscopy and proposed membrane topology of PfDMT1 **(A)** and yeast Smf3 **(B).** The intensity plots depict the normalized GFP fluorescence as a function of distance along the white line. The scissors in panel B represent FV proteases expected to cleave off a C-terminal GFP tag.

### PfDMT1 localizes to the FV membrane in an export-competent orientation

DMT1 homologs in humans and yeast commonly localize to the endosome/lysosome and vacuole membrane, respectively, where they mediate iron transport from the lumen of these compartments into the cytosol (24, 41). Two recent studies localized DMT1 from *P. falciparum* and *P. yoelli* to the FV membrane but did not specify orientation. To test and refine this localization in *P. falciparum*, we transfected Dd2 parasites with an episome encoding expression of PfDMT1 with a C-terminal GFP tag. Analysis of these parasites by epifluorescence microscopy revealed an annular GFP signal that encircled the hemozoin crystals, which were readily observed as dark pigments in brightfield images (Fig. 2A and Fig. S4A). Prior studies have shown that GFP tags exposed to the FV lumen are cleaved by resident proteases within this compartment, resulting in diffuse FV fluorescence (42, 43). Our observation of annular GFP fluorescence, therefore, suggests that the C-terminus of PfDMT1 extends into the cytosol (Fig. 2A). To test this conclusion, we transfected Dd2 parasites with an episome for expression of yeast Smf3 with a C-terminal GFP tag. Smf3 contains 11 transmembrane domains, which, based on Smf3 orientation on the yeast vacuole, would predict the C-terminal GFP would extend into the FV lumen opposite from the cytosolic N-terminus. In contrast to the ring-like fluorescence pattern of PfDMT1-GFP, we observed that Smf3-GFP exhibited diffuse signal throughout the FV (Fig. 2B and Fig. S4B) similar to FV-targeted matrix proteins (42, 43). The stable fluorescence of PfDMT1-GFP on the FV periphery strongly suggests that PfDMT1 is oriented in the FV membrane such that both the N- and C-termini face the cytosol. This PfDMT1 topology parallels the human DMT1 orientation on the endosomal/lysosomal membrane and is consistent with iron export from the FV lumen (Fig. 1B) (44).

### PfDMT1 is essential for parasite viability

To directly test if PfDMT1 is essential for blood-stage parasites, we used CRISPR/Cas9 to edit the PfDMT1 gene in NF54 PfMev parasites (45) to encode a C-terminal 5x hemagglutinin (5HA) tag and the aptamer/TetR-DOZI system for inducible knockdown (KD) (46, 47). In this system, protein expression is controlled by the non-toxic small molecule anhydrotetracycline (aTC) such that protein expression occurs normally in +aTC conditions but is stringently repressed in -aTC conditions (Fig. S5A and B). Correct genomic integration was confirmed by PCR (Fig. S5C and S5D).

Although PfDMT1-5HA has a predicted molecular mass ∼84 kDa, we detected a dominant band ∼55 kDa that was substantially lower than the expected size (Fig. 3A and Fig S6, A and B). This anomalous migration may be due to the presence of 12 hydrophobic TMDs that alter intrinsic electrophoretic mobility and/or to proteolytic processing of the lengthy 225-residue N-terminal sequence that precedes the first TMD and includes a span of 32 Asn residues. Removal of most or all of this N-terminal sequence would decrease the molecular mass of PfDMT1 by ∼25 kDa. We also observed an anomalous migration by SDS-PAGE/western blot for episomal PfDMT1-GFP, which has a predicted molecular mass of ∼105 kDa but migrated closer to 85 kDa (Fig. S6B). DMT1 homologs from other organisms have also been reported to migrate lower than expected, possibly due to their hydrophobic nature and/or proteolytic processing (35, 48). We observed normal PfDMT1 expression and parasite growth in +aTC conditions.

**Figure 3.**
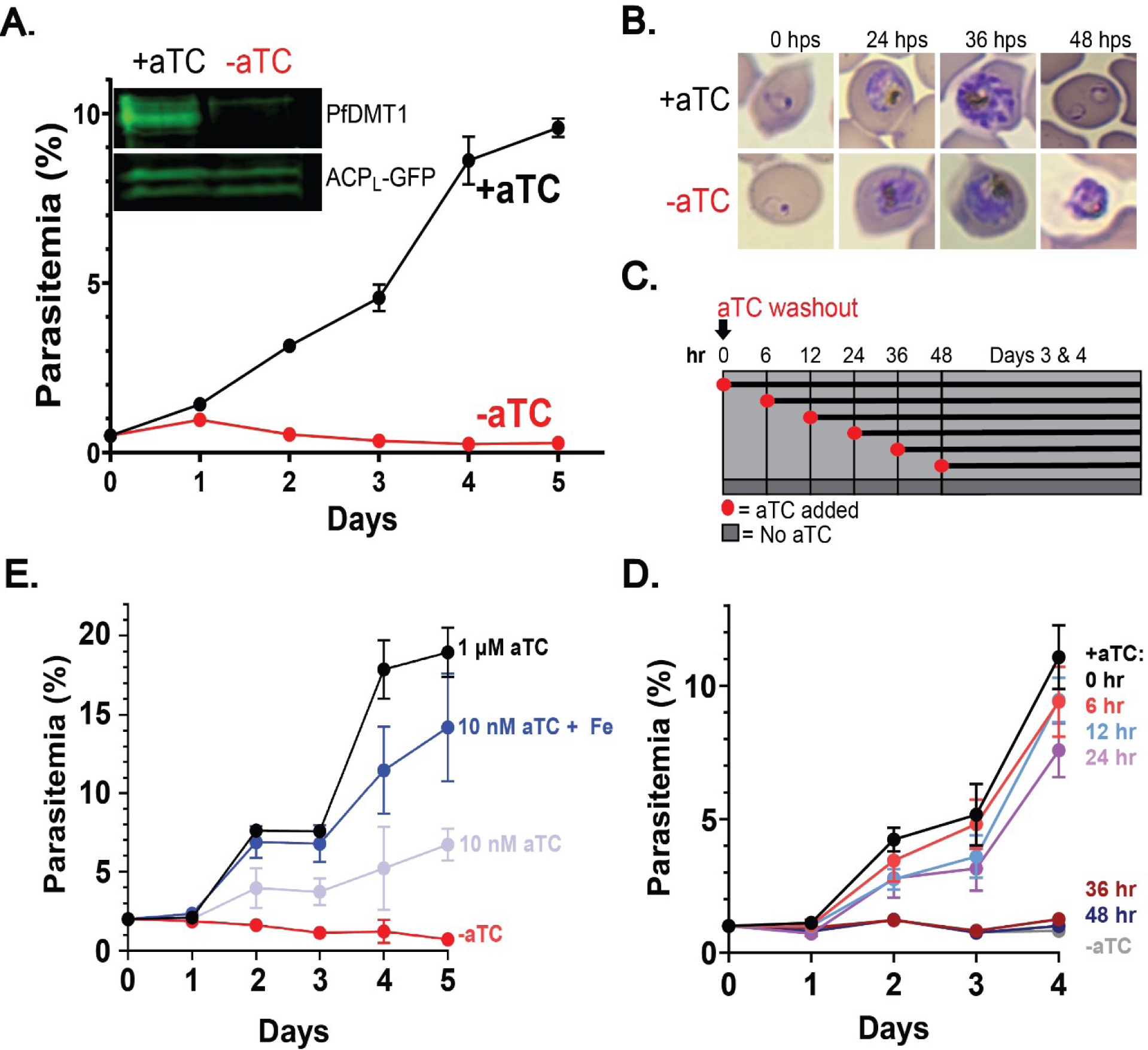
PfDMT1 is required for *P. falciparum* growth in the RBC and labile iron utilization. **(A)** Continuous growth assay of PfDMT1-5HA-aptamer-TetRDOZI parasites. Parasitemia is the percentage of RBCs infected with a parasite. Inset is a WB of PfDMT1-5HA and ACP_L_-GFP in parasites cultured ±aTC. **(B)** Giemsa-stained blood smears of *P. falciparum* grown ±aTC at listed hours post synchronization (hps). **(C)** Scheme of delayed aTC rescue of PfDMT1 KD. Parasites were synchronized (4-hour window) and treated with 1 µM aTC at indicated times. **(D)** Growth assay of delayed aTC rescue of PfDMT1 KD. **(E)** Continuous growth assay of PfDMT1 KD parasites grown ±10 nM or 1 µM aTC ± 125 µM FeCl_2_. Growth assay data points represent the mean ± SD of 6-9 biological replicates. We also tested iron supplementation at 5 nM and 2 nM aTC concentrations (Fig. S13).

However, PfDMT1 expression was undetectable in -aTC conditions, which resulted in rapid parasite death (Fig. 3A and Fig. S7). In our prior studies using the aptamer/TetR-DOZI system (13, 49, 50), growth impairment due to KD of essential proteins is usually observed in the second RBC-infection cycle after aTC washout. In contrast to these prior examples, PfDMT1 KD resulted in unusually fast parasite death that occurred in the same cycle as aTC washout, which was also reported for KD of PfEXP1 (51). To confirm the timing of parasite death, we seeded a growth assay at 3% parasitemia which provided greater separation of growth values between ±aTC conditions and confirmed that parasites failed to expand into a second infection cycle (Fig. S8). Blood-smear analysis revealed that -aTC parasites did not advance beyond the trophozoite stage. By 48 hours post-synchronization, stalled trophozoites could be observed inside and outside RBCs (Fig. 3B). These findings establish that PfDMT1 is critical for the survival and growth of blood-stage *P. falciparum* and strongly suggest a dominant functional contribution to metal transport at the FV.

To dissect the time course of parasite viability after aTC removal, we washed out aTC in highly synchronous (4-hour window) ring-stage parasites, reintroduced aTC at different time intervals, and measured parasite growth for several days (Fig. 3C). While parasites grew normally if aTC was added within 24 hours of removal, withholding aTC for 36 hours or longer resulted in irreversible parasite growth inhibition (Fig. 3D). This time-course was phenotypically similar to that observed for treatment with the iron chelator deferoxamine (DFO), which also kills *P. falciparum* with first-cycle kinetics (52, 53). Parasites were rescued from DFO treatment by adding exogenous iron to culture media and grew indistinguishably if iron was added within 24 hours of initiating DFO treatment (Fig. S9A and Fig. S9B). Delaying iron supplementation for 36 hours or longer, however, resulted in substantial growth inhibition. This phenotypic similarity is consistent with a model whereby PfDMT1 function is required for iron acquisition by blood-stage parasites.

Although parasites do not require exogenous iron in culture media for viability and growth, we posited that exogenous iron might rescue parasites from loss of PfDMT1 expression by increasing bioavailable iron within the infected RBC. However, exogenous supplementation with non-toxic concentrations of iron or other metals did not rescue parasites from PfDMT1 KD (Fig. S10A and Fig. S10B). Based on the functional model that parasites generate labile iron within the FV and depend on PfDMT1 transport to access this iron for metabolic utilization, we hypothesized that stringent KD of PfDMT1 expression might render FV iron inaccessible to parasites. This model predicted that less stringent, partial KD of PfDMT1 might result in residual PfDMT1 expression and a less severe growth phenotype that could be rescued by exogenous metal supplementation transported by residual PfDMT1.

To test this model, we first determined the dose-response curve for parasite growth at variable aTC concentrations and obtained an EC_50_ value of 7.5 nM aTC (Fig. S11), consistent with prior aTC EC_50_ values reported using the aptamer/TetR-DOZI system (54). Continuous growth assays performed at 10 nM aTC revealed substantial parasite growth inhibition compared to fully permissive 1 µM aTC growth conditions (Fig. 3E). Culture supplementation with iron almost entirely rescued parasite growth at 10 nM aTC to parasitemia values observed at 1 µM aTC, supporting the model that PfDMT1 function is necessary for iron acquisition (Fig. 3D). In contrast to strong rescue by Fe^2+^, supplementation with non-toxic concentrations of Ca^2+^, Zn^2+^, Mg^2+^, or Mn^2+^ resulted in minimal or no rescue of parasite growth at 10 nM aTC (Fig. S12), suggesting a specific and functionally dominant role for PfDMT1 in the transport of iron. Based on these observations, we conclude that PfDMT1 function is essential for blood-stage *P. falciparum* viability by gating access to labile iron in the FV.

### PfDMT1 KD impairs apicoplast biogenesis and mitochondrial polarization

Our model that PfDMT1 exports labile iron from the FV to support broad cellular metabolism predicted that PfDMT1 KD would impair iron-dependent processes outside the FV that occur in multiple sub-cellular compartments (Fig. 4A). To test this prediction, we studied the impact of PfDMT1 KD on biogenesis of the apicoplast organelle and the mitochondrial electron transport chain. These functions depend on iron but have not previously been linked to the FV. We first examined apicoplast biogenesis, which initiates early in schizogony and involves sequential elongation and branching before segmentation into discrete organelles that are partitioned with daughter merozoites (55). Prior work showed that DFO treatment substantially impairs apicoplast elongation, providing evidence that this process depends on iron (53). Subsequent studies also established that the apicoplast SUF Fe-S cluster synthesis pathway is needed to produce functional iron metalloproteins (IspG, IspH, and ferredoxin) that are required for apicoplast biogenesis through isoprenoid precursor synthesis (50, 56).

**Figure 4.**
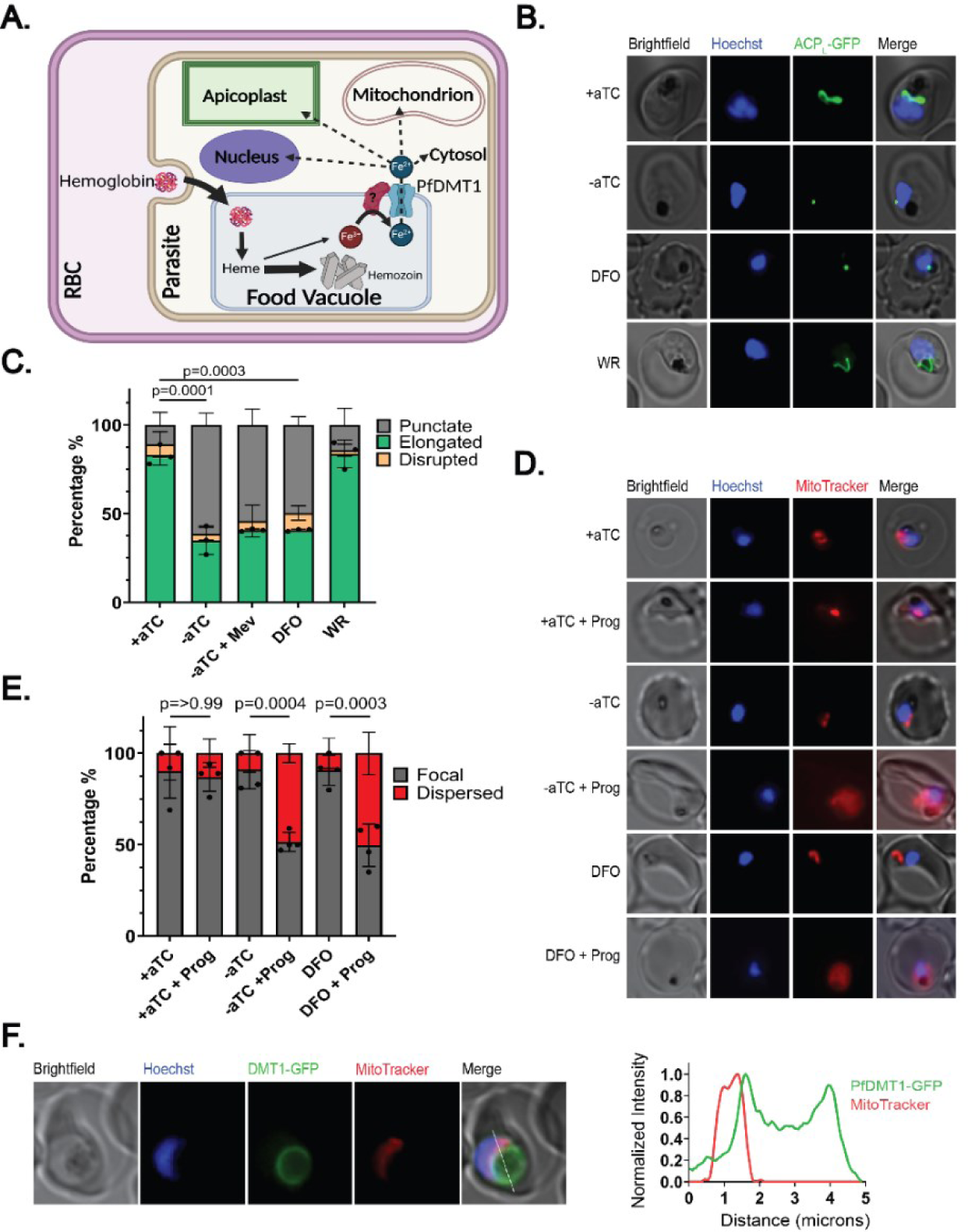
PfDMT1 KD impairs apicoplast and mitochondrial functions. **(A)** Proposed model of iron export via PfDMT1 to support broad cellular metabolism (figure created using BioRender.com). **(B)** Live-cell images of PfDMT1 KD parasites treated ± aTC, 60 µM DFO, 50 µM DL-mevalonate (Mev) or 5 nM WR. The apicoplast was visualized by ACP_L_-GFP expressed in PfMev NF54 parasites. **(C)** Population analysis of apicoplast morphology in each experimental condition. **(D)** Live-cell images of PfDMT1 KD parasites stained with MitoTracker™ Red CMXRos after culture ± aTC or treatment with 60 µM DFO ±5 µM proguanil (added 12 hours before imaging). **(E)** Population analysis of mitochondrial polarization in each experimental condition. For (C) and (E) each data point is the mean of biological replicates where 12-20 parasites were imaged and scored. A two-tailed unpaired t-test was utilized to calculate the given p-values (full comparison given in Table S1 and S2). In all experiments, parasites were synchronized to a 4-hour window and treated at t=0 hours post-synchronization to rings. **(F)** Live-cell image of PfDMT1-GFP and MitoTracker™ Red CMXRos signals. The intensity plot depicts the normalized GFP and MitoTracker fluorescence as a function of distance along the white line.

Using the PfDMT1 KD line in PfMev NF54 parasites (which express ACP_L_-GFP as a biomarker of apicoplast morphology)(45), we first confirmed that a lethal dose of DFO (60 µM) but not WR99210 (5 nM, cytoplasmic DHFR inhibitor) substantially blocked apicoplast elongation (Fig. 4, B and C), supporting the conclusion that one or more steps in apicoplast biogenesis specifically require labile iron. We also used fosmidomycin (FOS) as a positive control for the inhibition of apicoplast branching (Fig. S14A)(50). We then tested the impact of PfDMT1 KD in -aTC conditions and observed a strong impairment of apicoplast elongation that phenocopied DFO and FOS (Fig. 4 B and C). Both DFO treatment and PfDMT1 KD also blocked nuclear division such that parasites displayed a single, discrete focus stained by Hoescht (Fig. 4B). We considered the possibility that impaired apicoplast elongation from these treatments might be an indirect effect of impaired progression into schizogony. We noted, however, that WR99210 treatment also impaired nuclear division but did not block apicoplast elongation (Fig. 4B), providing evidence that initiation of apicoplast biogenesis is independent of nuclear division. We conclude that the impairment of apicoplast elongation is a specific phenotype of PfDMT1, supporting the model that labile iron required for apicoplast biogenesis is exported from the FV via PfDMT1 (Fig. 4A.).

The metabolic dependence of apicoplast biogenesis on iron is incompletely defined but includes a critical role for Fe-S clusters uniquely required to support isoprenoid precursor synthesis (56, 57). To test if loss of isoprenoid synthesis alone could account for defective apicoplast biogenesis upon PfDMT1 KD, we added exogenous mevalonate to the PfMev NF54 parasites into which PfDMT1 KD had been engineered. These parasites contain an engineered cytoplasmic pathway to synthesize isopentenyl pyrophosphate from mevalonate (45). The addition of mevalonate in -aTC conditions provided no rescue of apicoplast biogenesis. This observation suggests that Fe-dependent processes beyond Fe-S cluster-dependent IPP synthesis contribute to apicoplast biogenesis (56, 57).

*P. falciparum* parasites polarize the inner mitochondrial membrane via two redundant mechanisms that include proton translocation by complexes III and IV of the electron transport chain (ETC) and an undefined second mechanism that is inhibited by the drug proguanil (13, 58, 59). Proton pumping by the ETC depends on iron, due to essential contributions by the Rieske Fe-S protein to Complex III and by a parasite-specific Fe-S protein (ApiCox13) to Complex IV (60). These ETC complexes also require heme-dependent cytochromes (13), but blood-stage *Plasmodium* can scavenge host-derived heme and does not rely on de novo iron-dependent heme synthesis (14, 16, 61). ETC disruption is lethal to parasites due to impaired ubiquinone recycling but does not depolarize the mitochondrion due to redundant contributions by the second polarization mechanism. However, when the mechanism is blocked by proguanil, dysfunction of the ETC sensitizes parasites to mitochondrial depolarization such that positively charged mitochondrial dyes like MitoTracker fail to accumulate in the organelle (58). Prior knockdown studies of cytochrome *c_1_* and Rieske in Complex III support this conclusion (13, 49, 62).

We predicted that PfDMT1 KD would restrict mitochondrial iron availability and thus impair Fe-S synthesis crucial for Rieske- and ApiCox13-dependent ETC function (Fig. 4A) and sensitize parasites to mitochondrial depolarization by proguanil. Our positive control was a combination of the ETC inhibitor atovaquone and proguanil, which has previously been shown to depolarize the parasite mitochondrion (Fig. S14B) (58). Loss of PfDMT1 alone in -aTC conditions did not depolarize mitochondria, which accumulated MitoTracker Red CMXRos in discrete fluorescent foci similar to +aTC conditions. However, combining PfDMT1 KD with 5 µM proguanil treatment resulted in dispersed MitoTracker staining indicative of depolarization in >50% of parasites. Notably, DFO treatment phenocopied the effects of PfDMT1 KD, showing dispersed MitoTracker staining when combined with proguanil (Fig. 4, D and E). Our results support the model that ETC function and mitochondrial polarization depend on labile iron exported from the FV by PfDMT1.

The iron-trafficking mechanisms that connect PfDMT1 to the apicoplast and mitochondrion remain undefined. However, MitoTracker Red and PfDMT1-GFP signals were consistently observed in close proximity (Fig. 4F and Fig. S4A), which suggests that iron may directly transit from PfDMT1 on the FV periphery to the mitochondrion without a cytoplasmic intermediate, analogous to what has been observed at transient endosome/lysosome-mitochondrial contact sites in mammalian cells (23, 63, 64). A similar mechanism may account for iron delivery to the apicoplast, which is also frequently observed in close proximity to the FV (65). We have ongoing studies to unravel the mechanisms that traffic iron from PfDMT1 to sites of cellular utilization.

### Artemisinin activity does not depend on PfDMT1

Artemisinin (ART) and its derivatives are used in combination with other drugs as frontline malaria treatments, especially in Africa where virulent *P. falciparum* parasites are responsible for most infections and mortality (66). ART is a prodrug whose endoperoxide group is reductively cleaved via interaction with a reduced iron source within parasites to unleash its antimalarial activity. Prior works have provided strong evidence that ART activation depends on hemoglobin import and digestion within the FV (67, 68). The consensus view is that labile heme rather than exchangeable iron is the key activating factor, although conflicting evidence has been published regarding the impact of iron chelators like DFO on ART activity against blood-stage parasites (69–71). A recent study of rodent infection by *P. yoelii* presented evidence that reduced PyDMT1 expression diminished parasite sensitivity to ART, suggesting that iron transport from the FV is a critical axis of drug activation (28). We set out to test this conclusion and its relevance for human infection by *P. falciparum*, especially as our data supported a model whereby labile iron is transported out of the FV by PfDMT1 and stringently blocked by robust PfDMT1 KD.

Because clinically relevant ART resistance is mediated by differential activity against ring-stage parasites, we used a ring-stage survival assay (RSA) (72) to test if loss of PfDMT1 altered parasite sensitivity to ART. PfDMT1 KD schizonts were isolated by magnet purification and allowed to reinvade in fresh RBCs for 4 hours before sorbitol synchronization. To ensure stringent KD of PfDMT1 expression in ring-stage parasites, aTC was washed out of schizont-stage parasites 42 hours post synchronization. Washout of aTC in this fashion did not alter schizont development or diminish parasite ability to rupture and reinvade new RBCs compared to +aTC conditions (Fig. S15). These schizonts were then allowed to reinvade new RBCs for 6 hours after aTC washout. These parasites were then sorbitol synchronized, split into ±aTC growth conditions, and exposed to 700 nM dihydroartemisinin (DHA) for six hours (Fig. 5A). After DHA washout, all parasites were cultured +aTC to restore essential PfDMT1-mediated transport and analyzed by flow cytometry to determine parasitemia 66 hours later relative to DHA-untreated comparison cultures. As a positive control, we also studied the Cambodian *P. falciparum* isolate MRA-1240 (73, 74), which contains the R539T mutation in the kelch-13 protein previously shown via RSA to enhance parasite survival of DHA treatment. Although ∼70% of R539T parasites survived 700 nM DHA treatment compared to untreated controls, <20% of PfDMT1 KD parasites survived this treatment compared to untreated cultures (Fig. 5B). There was no difference in the survival of PfDMT1 KD parasites cultured ±aTC. Despite peak PfDMT1 transcription in ring-stage parasites (75, 76), we find no evidence that stringent PfDMT1 KD alters the ring-stage sensitivity of *P. falciparum* to DHA.

**Figure 5.**
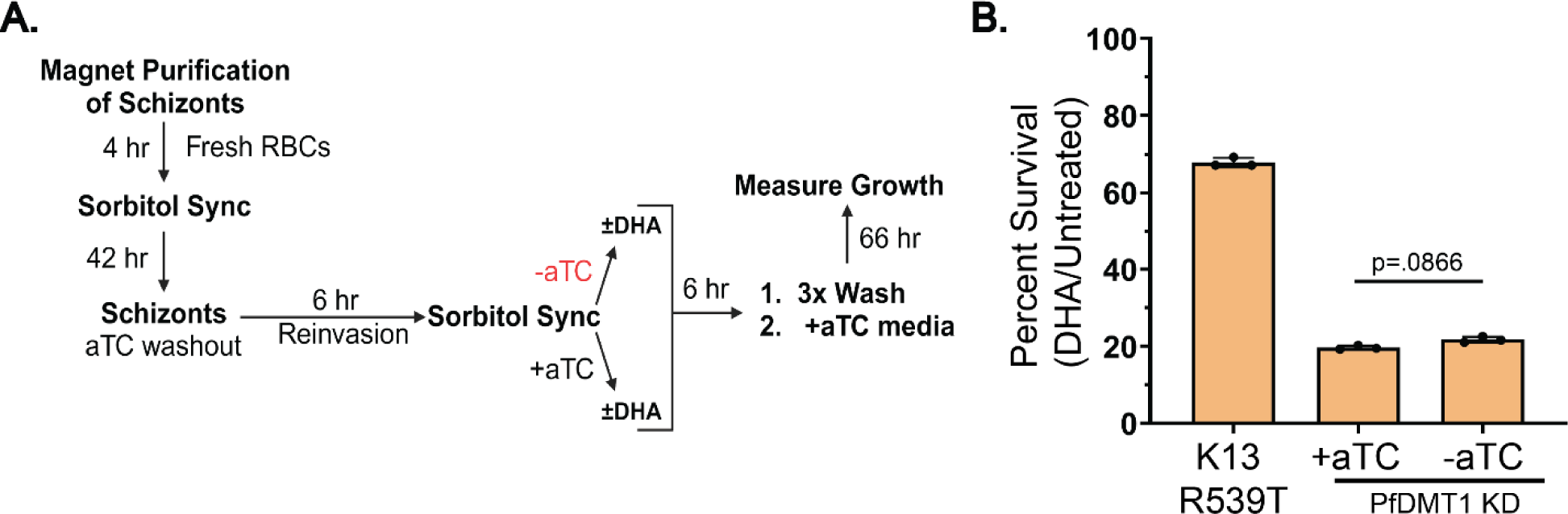
Loss of PfDMT1 does not alter parasite sensitivity to artemisinin in ring-stage parasites. **(A)** Schematic summary of ring-stage survival assay with 700 nM dihydroartemisinin (DHA) and PfDTM1 KD parasites cultured ±aTC. **(B)** Survival of ring-stage Cambodian isolate MRA-1240 with R539T K13 mutation and PfDMT1 KD parasites cultured ± aTC and pulsed with 700 nM DHA for 6 hours. Each data point represents the growth percentage of DHA-treated versus untreated parasites outgrown for 66 hours after DHA exposure. Percentages are the mean ± SD of biological triplicate assays analyzed by two-tailed unpaired t-test.

Because the FV is less prominent early in the lifecycle, we modified the RSA to allow PfDMT1 KD parasites to develop into trophozoites ±aTC before the 6-hour DHA pulse (Fig. S16A). Parasites grown in -aTC conditions displayed identical artemisinin sensitivity in the trophozoite stage relative to +aTC conditions (Fig. S16B). We conclude that PfDMT1 function and metal transport from the FV in blood-stage parasites do not contribute substantially to clinically relevant ART activation.

## DISCUSSION

During infection of human RBCs, *P. falciparum* malaria parasites utilize diverse mechanisms to obtain essential nutrients from the host cell and/or surrounding medium, including the acquisition of iron required by all cells for growth and viability. Although parasites infect the most iron-rich cells in the human body and are particularly sensitive to iron chelators, the fundamental mechanisms of iron acquisition and trafficking by blood-stage parasites have been unknown. This uncertainty has been a major blind spot in our basic understanding of parasite survival strategies within the host RBC. We have identified the acidic FV as a dominant source of labile iron for blood-stage *P. falciparum*, which utilizes the conserved iron transporter PfDMT1 to export FV iron for broad cellular utilization. This discovery unveils a key nutrient-acquisition mechanism in malaria parasites that can serve as a new therapeutic target and establishes a mechanistic foothold for unraveling cellular iron metabolism in this virulent pathogen.

### PfDMT1 function and specificity

Our results establish a cellular paradigm for blood-stage *P. falciparum* in which PfDMT1 functions at the FV membrane as the critical gatekeeper of labile iron generated within the FV (Fig. 4A). Parasites could be rescued by exogenous iron from partial but not stringent PfDMT1 knockdown (Fig. 4E), as expected if PfDMT1 is the dominant or exclusive mediator of essential iron acquisition during *Plasmodium* infection of RBCs. In this regard, PfDMT1 has favorable properties for therapeutic inhibition as an antimalarial drug target.

We consider trace non-enzymatic degradation of heme released by hemoglobin digestion to be the most likely source of exchangeable iron generated within the oxidizing FV environment, given the large-scale digestion of Hb and release of heme in this compartment (Fig. 4A). Nevertheless, import and digestion of other iron-containing RBC proteins (e.g., ferritin) may also contribute to exchangeable iron in the FV. This model of labile iron generation in the FV provides a unifying explanation for prior observations that iron chelators like DFO effectively restrict parasite growth but must be internalized into parasites to exert substantial anti-*Plasmodium* activity (6–9). Our observation that PfDMT1 knockdown has no measurable impact on ring-stage ART activation, which is sensitive to perturbations in Hb uptake and digestion (69, 70), suggests that PfDMT1 transport on the FV membrane functions independently from mechanisms of hemoglobin uptake and degradation.

We observed that parasite growth was strongly impaired in the same cell cycle in which PfDMT1 knockdown was initiated by aTC washout (Fig. 2A). This rapid sensitivity differs from our prior knockdown studies of proteins localized to the apicoplast and mitochondrion, in which growth phenotypes were not observed until at least the second cycle of RBC infection (13, 49, 50, 77). The unusual sensitivity of parasites to PfDMT1 knockdown may reflect the need to fully synthesize this protein de novo in each life cycle, since the FV is not inherited by daughter parasites (unlike the apicoplast and mitochondrion). We note that PfDMT1 transcription initiates and peaks in ring-stage parasites (75, 76), which supports this hypothesis. Nevertheless, parasites deprived of PfDMT1 expression remain viable for 24 hours and appear to developmentally transition to trophozoites before stalling their growth (Fig. 2, B and D). Sustained PfDMT1 knockdown beyond 24 hours causes irreversible parasite death, revealing a critical reliance on iron uptake for continued development of parasites into schizogony. These observations suggest that parasites inherit some labile iron and iron-containing proteins from the prior life cycle to satisfy initial metabolic requirements for ring-stage growth and/or that defects in iron-dependent metabolism do not critically limit blood-stage parasite growth until the trophozoite to schizont transition.

We observed that iron, but not other tested metals, rescued parasites from partial PfDMT1 knockdown. This observation suggests that iron is the functionally dominant substrate of PfDMT1 required by blood-stage parasites. As noted earlier, the Met-Ser divergence in the predicted metal-transport center of PfDMT1 is expected to impact its intrinsic metal specificity (78). Prior studies of DMT1 orthologs have suggested that the sulfur atom of Met provides a soft Lewis base that preferentially interacts with softer Lewis acids like Fe^2+^ or Mn^2+^ and facilitates their selective transport compared to harder Lewis acids like Ca^2+^ that are generally more abundant in the cell and which interact more favorably with hard bases like the oxygen atom of Ser (34). Our observation that iron is the critical substrate transported by PfDMT1 may suggest that there are additional amino-acid changes that tune metal ion specificity and/or that PfDMT1 also transports other metals that have additional, secondary mechanisms for uptake. We have initiated direct-transport assays to interrogate the metal specificity of PfDMT1 and the impact of this Met-Ser change. The prior report that *P. yoelii* DMT1, which also contains this mutation, functionally complements *S. cerevisiae* yeast deficient in Smf3 and rescues an iron-deficiency phenotype (28) suggests functional similarity in iron transport by yeast Smf3 and *P. yoelii* DMT1 despite this sequence difference. Nevertheless, this mutation along with other sequence differences between parasite and human DMT1 (Fig. 1A) may provide a molecular basis for developing selective inhibitors of PfDMT1 for antimalarial therapy or prophylaxis.

### Dependence of cellular iron metabolism on PfDMT1 and the FV

Our study provides direct evidence that key functions of the mitochondrion and apicoplast depend critically on PfDMT1 and iron obtained from the parasite FV. Although prior work has revealed that FV function requires apicoplast-derived isoprenoid precursors needed for prenylated Rab GTPases involved in vesicular trafficking of Hb to the FV (79, 80), a reciprocal dependence of the apicoplast on FV function has not been recognized. This functional interdependence further highlights that PfDMT1 operates as the linchpin in a critical axis of intracellular iron metabolism that connects essential functions across multiple compartments and organelles.

It remains uncertain if all or only a portion of intracellular iron needs depend on PfDMT1. Iron metabolism is compartmentalized in cells, and it is possible that only certain labile iron pools require PfDMT1 function to be filled. Nevertheless, we noted striking phenotypic similarity between PfDMT1 KD and treatment with DFO (Fig. 3, Fig. S9, and Fig. 4), which is expected to broadly chelate bioavailable iron within parasites. We, therefore, hypothesize that iron transport through PfDMT1 is the dominant or exclusive mechanism of iron acquisition by blood-stage *P. falciparum*. We are developing probes of iron-dependent metabolism in other compartments (e.g., cytoplasm and nucleus) to rigorously test this hypothesis.

### Phylogenetic conservation and divergence of DMT1 in Apicomplexa

DMT1 homologs are present in *Plasmodium* species and in *Toxoplasma gondii* but appear to be absent in other apicomplexan parasites, including *Babesia* and *Theileria.* Despite this common retention of DMT1, the *Plasmodium* and *T. gondii* homologs have notable differences in sequence and function. *T. gondii* and human DMT1 share 43% sequence identity that includes the conserved Met, compared to 29% sequence identity for *P. falciparum* and human DMT1 and Met-Ser mutation. In addition, DMT1 is essential for blood-stage *P. falciparum* but appears to be dispensable for *T. gondii* tachyzoites (81). The evolutionary path and fitness pressures that underpin retention and divergence of DMT1 in *T. gondii* and *Plasmodium* and loss in other apicomplexan parasites remain a key challenge to understand.

### PfDMT1 interaction partners and iron trafficking in *P. falciparum*

Cellular iron metabolism requires a network of interacting proteins that traffic, chaperone, and store bioavailable iron for utilization in discrete pathways and subcellular compartments. This key network has been difficult to unveil and understand in *Plasmodium* since parasites lack homologs of many canonical iron storage, trafficking, and transporter proteins found in well-studied yeast and mammalian cells (24, 81, 82). Identification of PfDMT1 as an essential mediator of labile iron in *P. falciparum* provides an exciting mechanistic foothold to unravel this network of protein interactors that have been refractory to identification by sequence analysis alone.

All DMT1 homologs specifically transport divalent metals and thus require a ferric reductase to convert Fe^3+^ to Fe^2+^ for transport (78). As such, PfDMT1 function is expected to require a ferric reductase partner to facilitate iron reduction in the oxidative FV environment, where iron is predominantly Fe^3+^. Parasites lack homologs of ferric reductases used in other organisms (e.g., STEAP3, Fre6) (24, 82), which represents a future challenge to fully understand *Plasmodium* iron mobilization from the FV. Additionally, iron exported from the FV by PfDMT1 is expected to be handed off to downstream trafficking proteins that are currently unknown (83). The proximity of PfDMT1 and the FV surface to the mitochondrion (Fig. 4F) and apicoplast (65) may suggest direct handoff to these organelles.

Iron importers into the mitochondrion and apicoplast are currently unknown, although a distant homolog of mammalian mitoferrin has been localized to the parasite mitochondrion (84). PfDMT1 topology in the FV membrane exposes the C-terminus to the cytoplasm (Fig. 2A) which will favor interaction studies by IP/MS and/or proximity biotinylation to identify key downstream client proteins that mediate iron uptake into organelles and trafficking in the cytoplasm, nucleus, and other compartments. Unveiling the network of protein interactors that shepherd iron from the FV to sites of utilization within the cell will elucidate fundamental parasite strategies for distributing this crucial nutrient and identify potential new targets for therapeutic intervention.

## MATERIALS AND METHODS

### Cloning

Episomal protein expression was accomplished by cloning the *P. falciparum* DMT1 (PF3D7_0523800) and yeast Smf3 (Uniprot Q12078) genes into pTEOE(15) between the XhoI and AvrII sites in frame with a C-terminal GFP tag. The pTEOE vector places protein expression under the control of the HSP86 promoter and includes a human DHFR positive selection cassette. Each gene was PCR-amplified from Dd2 parasite cDNA (primers 1/2) or the YcP-Smf3-GFP plasmid (primers 3/4, Supplementary file 1) (gift from Dr. Diane Ward, University of Utah), respectively, via ligation-independent cloning with the NEBuilder® HiFi DNA Assembly Master Mix (NEB E2621S). Correct plasmid insert sequences were confirmed by Sanger sequencing (University of Utah DNA Sequencing Core) using primers 14,15, 17, and 19 (Supplementary file 1).

### CRISPR-Cas9 Genome Editing of the PfDMT1 Locus

CRISPR/Cas9-stimulated repair by double-crossover homologous recombination was used to tag the PfDMT1 gene (PF3D7_0523800) to encode a 5x hemagglutinin (HA) C-terminal tag and the 10X aptamer/TetR-DOZI system (46, 47), which allows for PfDMT1 expression to be regulated by the non-toxic small molecule anhydrotetracycline (aTC, Caymen Chemicals 10009542). Ligation-independent cloning was used to insert annealed DNA oligos (primers 10/11 and 12/13, Supplementary file 1) into a modified pAIO CRISPR/Cas9 vector where the BtgZI cut site was replaced by HindIII for enhanced ease of use (49, 85). Two gRNA pAIO plasmids with district gRNA sequences were used in combination for transfections (g1: ATGGTTGTGAAAGAAATAAA, g2: AGTATTGCTATACAACTTTG).

To tag the PfDMT1 gene for knockdown, homology flanks consisting of the 427 bp of the 3’ UTR of PF3D7_0523800 separated from the final 693 bp (excluding the stop codon) of the coding sequence by an AflII cut site were inserted into the donor pKD repair plasmid (46) by ligation-independent cloning. Before transfection, the pKD vector was linearized by AflII digestion performed overnight at 37°C. Plasmid insert sequences were confirmed by Sanger DNA sequencing.

### Parasite Culturing and Transfection

All experiments were performed using *P. falciparum* Dd2 (86) or PfMev NF54 (45) parasites, whose identities were validated based on expected drug resistance/susceptibility. Parasite cultures were shown to be mycoplasma-free by PCR test. All culturing was done in Roswell Park Memorial Institute medium (RPMI-1640, Thermo Fisher 23400021) supplemented with 2.5 g/L Albumax I Lipid-Rich BSA (Thermo Fisher 11020039), 15 mg/L hypoxanthine (Sigma H9636), 110 mg/L sodium pyruvate (Sigma P5280), 1.19 g/L HEPES (Sigma H4034), 2.52 g/L sodium bicarbonate (Sigma S5761), 2 g/L glucose (Sigma G7021), and 10 mg/L gentamicin (Invitrogen Life Technologies 15750060), as described previously (13, 50). Cultures were grown in 2% hematocrit human RBCs received from the University of Utah Hospital blood bank, at 37°C and 5% CO_2_. Parasites were synchronized to a 4-hour window by magnet-purifying schizonts and then incubated with uninfected erythrocytes for 4 hours before treatment with 5% D-sorbitol (Sigma S7900).

To create episomal expression parasite lines with the pTEOE vector, 50–100 µg of purified plasmid was suspended in 1 x cytomix (87) with uninfected RBCs (uRBC) and transfected by electroporation using a Bio-Rad Gene Pulser Xcell system (0.31 kV, 925 µF). The electroporated RBCs were then seeded with Dd2 parasite culture at ∼1% and allowed to recover for 48 hours before drug selection with 5 nM WR99210 (Jacobus Pharmaceuticals). Stable transfectants emerged in 2-6 weeks.

PfDMT1 knockdown parasite lines were created by incubating 75 µg of linearized pKD donor plasmid, 50 µg of each pAIO plasmid (gRNA 3&4), and uninfected RBCs in 1x cytomix. Parasites were electroporated as described above. The electroporated RBCs were then seeded with PfMev NF54 parasites at ∼1% parasitemia allowed to grow for 48 hours in the presence of 1 µM aTC. Parasites were positively selected with blasticidin-S in the presence of 1 µM aTC for one week and then grown in no drug media + 1 µM aTC. Parasites returning from transfection were positively selected in blasticidin-S for one week. This was repeated until stable blasticidin parasites were recovered, as described previously (46). Polyclonal parasites returning from transfection were genotyped by PCR using primer sets 1/2, 17/18, and 19/20 (Supplementary file 1). Clonal parasites from independent transfections were isolated by limiting dilution and PCR-genotyped using the same primer sets as above (Fig. S5C and Fig. S5D). We observed indistinguishable growth phenotypes with all clonal integrates and selected clone D9 for further analysis.

### Parasite Growth Assays

For PfDMT1 KD growth assays, aTC was washed out 5x after 5% D-sorbitol treatment. PfDMT1 KD parasites were then split into ± aTC conditions before adding media with the listed concentrations of aTC. Growth assays were seeded at ∼1-2% parasitemia and monitored over multiple lifecycles with daily media changes. For metal-rescue experiments, aqueous stock solutions of 100 mM FeCl_2_, 100 mM CaCl_2_, 100 mM MgCl_2_, 1 mM ZnCl_2_, or 100 uM MnCl_2_ were directly diluted into parasite cultures at 125 µM (Fe^2+^, Ca^2+^, Mg^2+^), 5 µM (Zn^2+^), or 20n nM (Mn^2+^) final concentration. For experiments using DFO, a 10 mM aqueous stock solution of deferoxamine mesylate (Sigma-Aldrich 138-14-7) was diluted directly into cultures at 60 µM final concentration. Parasitemia was measured each day using flow cytometry by diluting 20 µL of parasite culture into 180 µL of 1.0 µg/mL acridine orange (Invitrogen Life Technologies A3568) in phosphate-buffered saline pH 7.5 (PBS) and analyzing on a BD FACSCelesta system monitoring SSC-A, FSC-A, PE-A, FITC-A, and PerCP-Cy5-5-A channels. Parasitemia of each biological replicate was determined by averaging two technical replicates. The final parasitemia was then determined by averaging the parasitemia of 3 or more biological replicates and calculating the standard deviation. To determine the EC50 of aTC in PfDMT1 KD parasites, ring-stage parasites within a 4-hour synchrony window were diluted to 1% parasitemia and incubated with a range of aTC concentrations through serial twofold dilution. The results of three biological replicates were fit to a 4-parameter dose-response curve. All graphs were plotted using GraphPad Prism 9.0.

### Fluorescence Microscopy

To stain the nuclei, parasites were incubated in a 0.1 µg/mL Hoechst 33342 (Thermo Scientific Pierce 62249) solution for ∼15 minutes at room temperature. The apicoplast was visualized 38 hours after synchronization in PfMev NF54 using the ACP_L_-GFP biomarker expressed in this line(45). FOS and DFO were added directly to 1 µM aTC culture at 10 µM and 60 µM final concentrations, respectively. WR-treated parasites were cultured in 5 nM WR99210 final concentration and 1 µM aTC. All drug treatments were initiated immediately after synchronization. The mitochondrion was visualized 36 hours after synchronization by staining parasites in 25 nM MitoTracker Red CMXROS (Invitrogen Life Technologies M7512) for 30 minutes before washing 3x in PBS or media. Atovaquone (DMSO) (Caymen Chemicals 95233184) and DFO (aq) were added to cultures at a final concentration of 100 nM and 60 µM, respectively (final [DMSO] <0.3%). Proguanil was added to parasite cultures 12 hours before cellular imaging at a final concentration 5 µM. 45–60 parasites in each condition were scored across 3-4 independent experiments to evaluate apicoplast morphology (focal or elongated) or MitoTracker signal (focal or dispersed). The statistical significance was determined using a two-tailed unpaired t-test using GraphPad Prism 9.0. Images were taken on DIC/brightfield, DAPI, GFP, and RFP channels with an EVOS M5000 imaging system. ImageJ was used to process and analyze images. All image processing (e.g., brightness, contrast) was done on a linear scale.

### Western Blot

To evaluate expression levels of PfDMT1-5HA, synchronized ∼5% parasitemia parasites were grown in ±aTC conditions and harvested 36 hours after synchronization. Parasite samples were incubated in a buffer consisting of 8 M Urea, 10 mM Tris-HCl, and 5% SDS and added to 5x SDS samples buffer containing 5% fresh β-mercaptoethanol. Samples were heated at 37°C for 60 minutes, loaded on 12.5% polyacrylamide gels, run at 120 V in the Bio-Rad mini-PROTEAN electrophoresis system, and transferred to nitrocellulose membrane using a 100 V setting for one hour via the Bio-Rad wet-transfer system. The membranes were blocked in 5% nonfat milk/PBS for one hour at room temperature, followed by overnight incubation with the primary antibody at 4°C, and then with the secondary antibody for one hour at room temperature. HA-tagged proteins were probed with a 1:1000 dilution of Roche rat anti-HA monoclonal 3F10 primary (Sigma 11867423001) and a 1:5000 dilution of donkey anti-rat DyLight800 (Invitrogen Life Technologies SA5-10032). Membranes containing GFP-tagged proteins were probed with a 1:1000 of goat anti-GFP polyclonal antibody (Abcam ab5450) and a 1:5000 dilution of donkey anti-goat IRDye800CW (Licor 926-32214). *P. falciparum* HSP60 was used as a loading control in PfDMT1-GFP parasites using a rabbit HSP60 antibody (Novus NBP2-12734) at 1:1000 dilution and then probed with anti-rabbit IRDye680 (Licor 926-68023). The Licor Odyssey system was used to image all membranes, with linear image processing in all cases.

### Immunoprecipitation

For Dd2 parasites with endogenously tagged PfDMT1-5HA, immunoprecipitation was conducted with Pierce anti-HA magnetic beads (Thermo Scientific 88836). 75 mL PfDMT1-5HA or Dd2 parasite pellets were lysed in 1 mL 1% Triton (Sigma 9002931). Each pellet was sonicated using a Branson sonicator equipped with a microtip probe and rotated at 4°C for 1 hr. Clarified lysates were collected after centrifugation at 17,000xg for 10 min. The anti-HA magnetic beads (30 µL) were equilibrated in 500 μL of cold 1% Trition in 1× PBS, and beads were collected using a magnetic stand. 1 mL of parasite lysates supernatant were incubated with anti-HA beads for 1 hr at 4°C while rotating. Anti-HA beads were washed using 1% Trition in 1× PBS 3x. Proteins bound to the anti-HA beads were eluted with 20 μL of 8 M urea in 10 mM Tris-HCl (pH 6.8) and stored at –80°C until use.

### Structural modeling

Predicted structural models for *P. falciparum* DMT1 and human DMT1 were obtained from the AlphaFold Protein Structure Database (https://alphafold.ebi.ac.uk)(88). Protein models were visualized and analyzed using PyMOL.

### Alignment and Phylogenetic Analyses

The protein sequence of *P. falciparum* DMT1 was analyzed by PSI-BLAST(89) to identify homologs in *Toxoplasma gondii*, *Vitrella brassicaformis*, *Saccharomyces cerevisiae*, and *Homo sapiens*. All other sequences were obtained from a previous study that rigorously characterized the four groups of eukaryotic DMT1 transporters(40). Sequences were aligned by CLUSTAL OMEGA(90). The multisequence alignment file was then uploaded to the Interactive Tree of Life (ITOL) webserver to generate a 1000 bootstrap maximum-likelihood phylogenetic tree (91).

### Ring-Stage Survival Assay (RSA)

Parasites were synchronized within a 4-hour synchrony window, and the RSA was performed as previously described(72). Briefly, schizont-stage PfDMT1 KD or Cambodian K13 R539T parasites were purified by passage over a magnetic column and then allowed to invade fresh RBCs for 4 hours before sorbitol synchronization. After 42 hours of growth following sorbitol treatment, aTC was washed out of the media. Parasites were then allowed to reinvade fresh RBCs for 6 hours without aTC being present. After 6 hours, parasites were treated with 5% D-sorbitol treatment and then split into ±aTC conditions and further separated in ±700 nM DHA (Sigma-Aldrich D7439) treatment. After 6 hours of DHA treatment, parasites were washed 3x and all parasite treatments were grown in +aTC media. The Cambodian isolate MRA-1240(73), which contains the R589T mutation in the kelch-13 protein, was used as a positive control and pulsed with DHA as described above. The parasitemia between DHA-treated and untreated parasites was then compared 66 hours after DHA treatment (72 hours after reinvasion).

## ACKNOWLEDGMENTS

We thank Diane Ward and members of the Sigala lab for helpful discussions. This work was supported by NIH grant R35GM133764 (to PAS) and a pilot award (to PAS) from the Utah Center for Iron and Heme Disorders (supported by U54DK110858). KML was supported by NIH training grant T32TR004392. DNA synthesis, sequencing, fluorescence microscopy, and flow cytometry were performed using core facilities at the University of Utah.

## REFERENCES

1. Anonymous (2022) World Malaria Report 2022. (World Health Orginization).

2. A. Trampuz, M. Jereb, I. Muzlovic, R. M. Prabhu, Clinical review: Severe malaria. Crit Care 7, 315–323 (2003).

3. B. Balikagala et al., Evidence of Artemisinin-Resistant Malaria in Africa. N Engl J Med 385, 1163–1171 (2021).

4. C. C. Philpott, M. S. Ryu, Special delivery: distributing iron in the cytosol of mammalian cells. Front Pharmacol 5, 173 (2014).

5. M. Loyevsky et al., Chelation of iron within the erythrocytic Plasmodium falciparum parasite by iron chelators. Mol Biochem Parasitol 101, 43–59 (1999).

6. M. D. Scott, A. Ranz, F. A. Kuypers, B. H. Lubin, S. R. Meshnick, Parasite uptake of desferroxamine: a prerequisite for antimalarial activity. Br J Haematol 75, 598–602 (1990).

7. M. Loyevsky et al., The antimalarial action of desferal involves a direct access route to erythrocytic (Plasmodium falciparum) parasites. J Clin Invest 91, 218–224 (1993).

8. T. E. Peto, J. L. Thompson, A reappraisal of the effects of iron and desferrioxamine on the growth of Plasmodium falciparum ‘in vitro’: the unimportance of serum iron. Br J Haematol 63, 273–280 (1986).

9. R. G. Marvin et al., Fluxes in “free” and total zinc are essential for progression of intraerythrocytic stages of Plasmodium falciparum. Chem Biol 19, 731–741 (2012).

10. P. A. Sigala, J. R. Crowley, S. Hsieh, J. P. Henderson, D. E. Goldberg, Direct tests of enzymatic heme degradation by the malaria parasite Plasmodium falciparum. J Biol Chem 287, 37793–37807 (2012).

11. A. M. Blackwell et al., Malaria parasites require a divergent heme oxygenase for apicoplast gene expression and biogenesis. bioRxiv 10.1101/2024.05.30.596652, 2024.2005.2030.596652 (2024).

12. P. F. Scholl, A. K. Tripathi, D. J. Sullivan, Bioavailable iron and heme metabolism in Plasmodium falciparum. Curr Top Microbiol Immunol 295, 293–324 (2005).

13. T. J. Espino-Sanchez et al., Direct tests of cytochrome c and c(1) functions in the electron transport chain of malaria parasites. Proc Natl Acad Sci U S A 120, e2301047120 (2023).

14. H. Ke et al., The heme biosynthesis pathway is essential for Plasmodium falciparum development in mosquito stage but not in blood stages. J Biol Chem 289, 34827–34837 (2014).

15. P. A. Sigala, J. R. Crowley, J. P. Henderson, D. E. Goldberg, Deconvoluting heme biosynthesis to target blood-stage malaria parasites. Elife 4 (2015).

16. V. A. Nagaraj et al., Malaria parasite-synthesized heme is essential in the mosquito and liver stages and complements host heme in the blood stages of infection. PLoS Pathog 9, e1003522 (2013).

17. P. Loria, S. Miller, M. Foley, L. Tilley, Inhibition of the peroxidative degradation of haem as the basis of action of chloroquine and other quinoline antimalarials. Biochem J 339 (Pt 2), 363–370 (1999).

18. E. Nagababu, J. M. Rifkind, Reaction of hydrogen peroxide with ferrylhemoglobin: superoxide production and heme degradation. Biochemistry 39, 12503–12511 (2000).

19. H. Atamna, H. Ginsburg, Origin of reactive oxygen species in erythrocytes infected with Plasmodium falciparum. Mol Biochem Parasitol 61, 231–241 (1993).

20. A. H. Bryk, J. R. Wisniewski, Quantitative Analysis of Human Red Blood Cell Proteome. J Proteome Res 16, 2752–2761 (2017).

21. M. Cazzola et al., Biologic and clinical significance of red cell ferritin. Blood 62, 1078–1087 (1983).

22. G. Clarebout et al., Status of Plasmodium falciparum towards catalase. Br J Haematol 103, 52–59 (1998).

23. F. Rizzollo, S. More, P. Vangheluwe, P. Agostinis, The lysosome as a master regulator of iron metabolism. Trends Biochem Sci 46, 960–975 (2021).

24. L. Ramos-Alonso, A. M. Romero, M. T. Martinez-Pastor, S. Puig, Iron Regulatory Mechanisms in Saccharomyces cerevisiae. Front Microbiol 11, 582830 (2020).

25. T. Sahu et al., ZIPCO, a putative metal ion transporter, is crucial for Plasmodium liver-stage development. EMBO Mol Med 6, 1387–1397 (2014).

26. K. Slavic et al., A vacuolar iron-transporter homologue acts as a detoxifier in Plasmodium. Nat Commun 7, 10403 (2016).

27. J. S. Wichers et al., PMRT1, a Plasmodium-Specific Parasite Plasma Membrane Transporter, Is Essential for Asexual and Sexual Blood Stage Development. mBio 13, e0062322 (2022).

28. M. Zhong, B. Zhou, Plasmodium yoelii iron transporter PyDMT1 interacts with host ferritin and is required in full activity for malarial pathogenesis. BMC Biol 21, 279 (2023).

29. Y. Nevo, N. Nelson, The NRAMP family of metal-ion transporters. Biochim Biophys Acta 1763, 609–620 (2006).

30. B. Mackenzie, M. L. Ujwal, M. H. Chang, M. F. Romero, M. A. Hediger, Divalent metal-ion transporter DMT1 mediates both H+-coupled Fe2+ transport and uncoupled fluxes. Pflugers Arch 451, 544–558 (2006).

31. S. Lam-Yuk-Tseung, G. Govoni, J. Forbes, P. Gros, Iron transport by Nramp2/DMT1: pH regulation of transport by 2 histidines in transmembrane domain 6. Blood 101, 3699–3707 (2003).

32. M. Zhang et al., Uncovering the essential genes of the human malaria parasite Plasmodium falciparum by saturation mutagenesis. Science 360 (2018).

33. M. P. Mims, J. T. Prchal, Divalent metal transporter 1. Hematology 10, 339–345 (2005).

34. A. T. Bozzi et al., Conserved methionine dictates substrate preference in Nramp-family divalent metal transporters. Proc Natl Acad Sci U S A 113, 10310–10315 (2016).

35. A. T. Bozzi et al., Crystal Structure and Conformational Change Mechanism of a Bacterial Nramp-Family Divalent Metal Transporter. Structure 24, 2102–2114 (2016).

36. I. A. Ehrnstorfer, E. R. Geertsma, E. Pardon, J. Steyaert, R. Dutzler, Crystal structure of a SLC11 (NRAMP) transporter reveals the basis for transition-metal ion transport. Nat Struct Mol Biol 21, 990–996 (2014).

37. I. A. Ehrnstorfer, C. Manatschal, F. M. Arnold, J. Laederach, R. Dutzler, Structural and mechanistic basis of proton-coupled metal ion transport in the SLC11/NRAMP family. Nat Commun 8, 14033 (2017).

38. L. Kall, A. Krogh, E. L. Sonnhammer, Advantages of combined transmembrane topology and signal peptide prediction--the Phobius web server. Nucleic Acids Res 35, W429–432 (2007).

39. J. A. Robledo, P. Courville, M. F. Cellier, G. R. Vasta, Gene organization and expression of the divalent cation transporter Nramp in the protistan parasite Perkinsus marinus. J Parasitol 90, 1004–1014 (2004).

40. M. F. M. Cellier, Nramp: Deprive and conquer? Front Cell Dev Biol 10, 988866 (2022).

41. I. Yanatori, F. Kishi, DMT1 and iron transport. Free Radic Biol Med 133, 55–63 (2019).

42. M. Klemba, W. Beatty, I. Gluzman, D. E. Goldberg, Trafficking of plasmepsin II to the food vacuole of the malaria parasite Plasmodium falciparum. J Cell Biol 164, 47–56 (2004).

43. J. M. Matz et al., A lipocalin mediates unidirectional heme biomineralization in malaria parasites. Proc Natl Acad Sci U S A 117, 16546–16556 (2020).

44. M. Czachorowski, S. Lam-Yuk-Tseung, M. Cellier, P. Gros, Transmembrane topology of the mammalian Slc11a2 iron transporter. Biochemistry 48, 8422–8434 (2009).

45. R. P. Swift et al., A mevalonate bypass system facilitates elucidation of plastid biology in malaria parasites. PLoS Pathog 16, e1008316 (2020).

46. K. Rajaram, H. B. Liu, S. T. Prigge, Redesigned TetR-Aptamer System To Control Gene Expression in Plasmodium falciparum. mSphere 5 (2020).

47. S. M. Ganesan, A. Falla, S. J. Goldfless, A. S. Nasamu, J. C. Niles, Synthetic RNA-protein modules integrated with native translation mechanisms to control gene expression in malaria parasites. Nat Commun 7, 10727 (2016).

48. C. Ballesteros, J. F. Geary, C. D. Mackenzie, T. G. Geary, Characterization of Divalent Metal Transporter 1 (DMT1) in Brugia malayi suggests an intestinal-associated pathway for iron absorption. Int J Parasitol Drugs Drug Resist 8, 341–349 (2018).

49. S. Falekun et al., Divergent acyl carrier protein decouples mitochondrial Fe-S cluster biogenesis from fatty acid synthesis in malaria parasites. Elife 10 (2021).

50. M. Okada et al., Critical role for isoprenoids in apicoplast biogenesis by malaria parasites. Elife 11 (2022).

51. T. Nessel et al., EXP1 is required for organisation of EXP2 in the intraerythrocytic malaria parasite vacuole. Cell Microbiol 22, e13168 (2020).

52. G. F. Mabeza, M. Loyevsky, V. R. Gordeuk, G. Weiss, Iron chelation therapy for malaria: a review. Pharmacol Ther 81, 53–75 (1999).

53. M. Okada, P. Guo, S. A. Nalder, P. A. Sigala, Doxycycline has distinct apicoplast-specific mechanisms of antimalarial activity. Elife 9 (2020).

54. S. J. Goldfless, J. C. Wagner, J. C. Niles, Versatile control of Plasmodium falciparum gene expression with an inducible protein-RNA interaction. Nat Commun 5, 5329 (2014).

55. E. Yeh, J. L. DeRisi, Chemical rescue of malaria parasites lacking an apicoplast defines organelle function in blood-stage Plasmodium falciparum. PLoS Biol 9, e1001138 (2011).

56. J. E. Gisselberg, T. A. Dellibovi-Ragheb, K. A. Matthews, G. Bosch, S. T. Prigge, The suf iron-sulfur cluster synthesis pathway is required for apicoplast maintenance in malaria parasites. PLoS Pathog 9, e1003655 (2013).

57. M. Okada, P. A. Sigala, The interdependence of isoprenoid synthesis and apicoplast biogenesis in malaria parasites. PLoS Pathog 19, e1011713 (2023).

58. H. J. Painter, J. M. Morrisey, M. W. Mather, A. B. Vaidya, Specific role of mitochondrial electron transport in blood-stage Plasmodium falciparum. Nature 446, 88–91 (2007).

59. T. S. Skinner-Adams et al., Cyclization-blocked proguanil as a strategy to improve the antimalarial activity of atovaquone. Commun Biol 2, 166 (2019).

60. F. Evers et al., Composition and stage dynamics of mitochondrial complexes in Plasmodium falciparum. Nat Commun 12, 3820 (2021).

61. D. E. Goldberg, P. A. Sigala, Plasmodium heme biosynthesis: To be or not to be essential? PLoS Pathog 13, e1006511 (2017).

62. P. Sheokand, A. Mühleip, L. Sheiner, *PLASMODIUM falciparum* mitochondrial complex III, the target of atovaquone, is essential for progression to the transmissible sexual stages. bioRxiv 10.1101/2024.01.09.574740, 2024.2001.2009.574740 (2024).

63. A. Hamdi et al., Erythroid cell mitochondria receive endosomal iron by a “kiss-and-run” mechanism. Biochim Biophys Acta 1863, 2859–2867 (2016).

64. J. Barra et al., DMT1-dependent endosome-mitochondria interactions regulate mitochondrial iron translocation and metastatic outgrowth. Oncogene 43, 650–667 (2024).

65. L. H. Bannister, J. M. Hopkins, R. E. Fowler, S. Krishna, G. H. Mitchell, A brief illustrated guide to the ultrastructure of Plasmodium falciparum asexual blood stages. Parasitol Today 16, 427–433 (2000).

66. E. G. Tse, M. Korsik, M. H. Todd, The past, present and future of anti-malarial medicines. Malar J 18, 93 (2019).

67. N. Klonis et al., Artemisinin activity against Plasmodium falciparum requires hemoglobin uptake and digestion. Proc Natl Acad Sci U S A 108, 11405–11410 (2011).

68. J. Birnbaum et al., A Kelch13-defined endocytosis pathway mediates artemisinin resistance in malaria parasites. Science 367, 51–59 (2020).

69. N. Klonis, D. J. Creek, L. Tilley, Iron and heme metabolism in Plasmodium falciparum and the mechanism of action of artemisinins. Curr Opin Microbiol 16, 722–727 (2013).

70. K. E. Ward, D. A. Fidock, J. L. Bridgford, Plasmodium falciparum resistance to artemisinin-based combination therapies. Curr Opin Microbiol 69, 102193 (2022).

71. J. Wang et al., Haem-activated promiscuous targeting of artemisinin in Plasmodium falciparum. Nat Commun 6, 10111 (2015).

72. C. Amaratunga, A. T. Neal, R. M. Fairhurst, Flow cytometry-based analysis of artemisinin-resistant Plasmodium falciparum in the ring-stage survival assay. Antimicrob Agents Chemother 58, 4938–4940 (2014).

73. B. Witkowski et al., Novel phenotypic assays for the detection of artemisinin-resistant Plasmodium falciparum malaria in Cambodia: in-vitro and ex-vivo drug-response studies. Lancet Infect Dis 13, 1043–1049 (2013).

74. F. Ariey et al., A molecular marker of artemisinin-resistant Plasmodium falciparum malaria. Nature 505, 50–55 (2014).

75. A. K. Subudhi et al., Malaria parasites regulate intra-erythrocytic development duration via serpentine receptor 10 to coordinate with host rhythms. Nat Commun 11, 2763 (2020).

76. M. J. Lopez-Barragan et al., Directional gene expression and antisense transcripts in sexual and asexual stages of Plasmodium falciparum. BMC Genomics 12, 587 (2011).

77. A. E. Garcia-Guerrero, R. G. Marvin, A. M. Blackwell, P. A. Sigala, Biogenesis of cytochromes c and c(1) in the electron transport chain of malaria parasites. bioRxiv 10.1101/2024.02.01.575742 (2024).

78. C. Manatschal, R. Dutzler, The Structural Basis for Metal Ion Transport in the SLC11/NRAMP Family. Chimia (Aarau) 76, 1005–1010 (2022).

79. K. Kennedy et al., Delayed death in the malaria parasite Plasmodium falciparum is caused by disruption of prenylation-dependent intracellular trafficking. PLoS Biol 17, e3000376 (2019).

80. R. Howe, M. Kelly, J. Jimah, D. Hodge, A. R. Odom, Isoprenoid biosynthesis inhibition disrupts Rab5 localization and food vacuolar integrity in Plasmodium falciparum. Eukaryot Cell 12, 215–223 (2013).

81. M. A. Sloan, D. Aghabi, C. R. Harding, Orchestrating a heist: uptake and storage of metals by apicomplexan parasites. Microbiology (Reading) 167 (2021).

82. D. M. Ward, S. M. Cloonan, Mitochondrial Iron in Human Health and Disease. Annu Rev Physiol 81, 453–482 (2019).

83. C. C. Philpott et al., Iron-tracking strategies: Chaperones capture iron in the cytosolic labile iron pool. Front Mol Biosci 10, 1127690 (2023).

84. J. Wunderlich et al., Iron transport pathways in the human malaria parasite *Plasmodium falciparum* revealed by RNA-sequencing. bioRxiv 10.1101/2024.04.18.590068, 2024.2004.2018.590068 (2024).

85. N. J. Spillman, J. R. Beck, S. M. Ganesan, J. C. Niles, D. E. Goldberg, The chaperonin TRiC forms an oligomeric complex in the malaria parasite cytosol. Cell Microbiol 19 (2017).

86. T. E. Wellems et al., Chloroquine resistance not linked to mdr-like genes in a Plasmodium falciparum cross. Nature 345, 253–255 (1990).

87. T. Ponnudurai, A. D. Leeuwenberg, J. H. Meuwissen, Chloroquine sensitivity of isolates of Plasmodium falciparum adapted to in vitro culture. Trop Geogr Med 33, 50–54 (1981).

88. J. Jumper et al., Highly accurate protein structure prediction with AlphaFold. Nature 596, 583–589 (2021).

89. S. F. Altschul et al., Gapped BLAST and PSI-BLAST: a new generation of protein database search programs. Nucleic Acids Res 25, 3389–3402 (1997).

90. F. Madeira et al., Search and sequence analysis tools services from EMBL-EBI in 2022. Nucleic Acids Res 50, W276–W279 (2022).

91. I. Letunic, P. Bork, Interactive Tree Of Life (iTOL) v5: an online tool for phylogenetic tree display and annotation. Nucleic Acids Res 49, W293–W296 (2021).

