## Supplementary Figures for "Identification of a divalent metal transporter required for cellular iron metabolism in malaria parasites"

<u>SUPPORTING FIGURES:</u>

```
P.falciparum       MEKDFTERSTKGQRKLANGVVENIESKLKNHLDNNNVINKSVNYEDVMCSNVKLSVHDDE  60
H.sapiens          ------------------------------------------------------------  0
S.cerevisiae       ------------------------------------------------------------  0

P.falciparum       YDIKNIKKDVCNLSKLSEERLSNNISLKDNNKVVKNPEIDTKTARQQYKKRLNSTQSTMN  120
H.sapiens          ------------------------------------------------------------  0
S.cerevisiae       ------------------------------------------------------------  0

P.falciparum       IEGNINNEKDDIIEDYDDGLIRNMDYEMSIVKNKRNNNNNNNNNNNNNNNNNNNNNNNNN  180
H.sapiens          ---------------------------MVLGPEQKMSDDSVSGDHGES---------ASL  24
S.cerevisiae       ------------------------------------------------------------  0

P.falciparum       NNNNNNNSNNVDNRKRNKNEECYDESYCISDNLDEITSYRNKLSLYNKLRMCFNYFGPGW  240
H.sapiens          GNINPAYSNPSLSQSPGDSEEYFATYF------NEKISIPEEEYSCFSFRKLWAFTGPGF  78
S.cerevisiae       -----------------------------------------MRSYMQILQKFAKFIGPGI  19
                                                                   ::    : ***

P.falciparum       IVAIAYLDPGNLCSNLNVGLIRSPDPTLEKDYSGYYLLWIMVYGHMLGFIFQVLSMRLGH  300
H.sapiens          LMSIAYLDPGNIESDLQSGA-----------VAGFKLLWILLLATLVGLLLQRLAARLGV  127
S.cerevisiae       LVSVAYMDPGNYATSVSGGA-----------QYKYTLLFSIFISNIFAVLLQCLCVKLGT  68
                   ::::**:****  :.:. *               : **: :. . :...::* *. :**

P.falciparum       VTGLDLASLCSKEFDRTTSTIIYVLVQIAIWGAHIQAIIGTFIALNLIF--GISVKVAIF  358
H.sapiens          VTGLHLAEVCHRQYPKVPRVILWLMVELAIIGSDMQEVIGSAIAINLLSVGRIPLWGGVL  187
S.cerevisiae       ITGYDLAENCRHNLPKKLNYTLYLFAEVAIIATDLAEVVGTAIALQILF--KIPLTWGVL  126
                   :** .**. * ::  :     ::::.::** .:.:  ::*: **::::    * :  .::

P.falciparum       YTLFEAIIYSFLENKSLGLL------ENVLSFLVGILAVSFFVNVFMTPINFKELAISIL  412
H.sapiens          ITIADTFVFLFL-DKYG--LRKLEAF---FGFLITIMALTFGYEYVT-VKPSQSQVLKGM  240
S.cerevisiae       LTVLDVLVILMFYTPNGQSLKKVRVFEFGVGILVIGTCICFVLELFKVSIPDKAELFKGF  186
                    *: :.::  ::       *         ..:*:   .: *  : .      :   :. :

P.falciparum       YPRIPKGK----EIDALALLGSIISAHIFYLHTNLTAKKKSVICNDLSLR-------RYN  461
H.sapiens          FVPSCSGCRTPQIEQAVGIVGAVIMPHNMYLHSALVKSRQVNRNNKQEVR-------EAN  293
S.cerevisiae       LPSNIIFKEQQALYISLGILGATVMPHSLYLGSSIVKPRLHDYDLKKYGKVNARPSLSAI  246
                                  ::.::*: :  * :** : :.  :      .   :

P.falciparum       TLGTIESGGSLFLSCLTNCIIVLTFAEVNL----------------------KSFERRDQ  499
H.sapiens          KYFFIESCIALFVSFIINVFVVSVFAEAFFGKTNEQVVEVCTNTSSPHAGLFPKDNSTLA  353
S.cerevisiae       KYTLNY----AYAELIISLFLIATFVNS-----AILIVAGATLSGQ------PEAED---  288
                   .          : . : . ::: .*.:                          . :

P.falciparum       YNLFTAYEVMRKSFGKISMYIWSFGLLSSGNNSSFMCEYASKSVVEGFLNKKINTFVRVF  559
H.sapiens          VDIYKGGVVLGCYFGPAALYIWAVGILAAGQSSTMTGTYSGQFVMEGFLNLKWSRFARVV  413
S.cerevisiae       ADLLSIYKLLVHYISPAAGLIFALAMLCSGQSAGIICTLAGQIVSEGFLQWSLPPWATRL  348
                    :: .   ::   :.  :  *::..:*.:*:.: :    :.: * ****: .   :.  .

P.falciparum       TFRLM-LFSLLYMFLTLNKYTLDQLTNFINVIQVLLLPMATIPLYRFSIHENVLGEFRLK  618
H.sapiens          LTRSIAIIPTLLVAVFQDVEHLTGMNDFLNVLQSLQLPFALIPILTFTSLRPVMSDFANG  473
S.cerevisiae       CTRLIAIVPCLFVTLTMGEKGISDILNFSQVVLSLILPIVSAPLIYFTANRKLMVVHDEN  408
                     * : :.  * : :  .   :  : :* :*:  * **:.  *:  *:  . ::  .

P.falciparum       KFPKFA-----------------------------------VFLIIIAIIISNVLLTFLD  643
H.sapiens          LGWRIAGGI--------------------------------LVLIICSINMYFVVVYVRD  501
S.cerevisiae       GVVRAPADVNAIADETTPLNSKHSKIVDFTNSRLLTYTSVFVWALIGSLNCYLVISYLLG  468
                      :                                     :  :* ::    *:  . .

P.falciparum       FVHKETSLITIFFLVIFSFLYFGFIIYFFN---IPIKKNYIQRN----------------  684
H.sapiens          LGHVA----LYVVAAVVSVAYLGFVFYLGWQCLIALGMSFLDCGHTCHLGLTAQPELYLL  557
S.cerevisiae       -----------------------------------ADIHF--------------------  473
                                                          :

P.falciparum       -----------   684
H.sapiens          NTMDADSLVSR   568
S.cerevisiae       -----------   473
```

**Figure S1**. **Sequence alignment of DMT1 homologs.** Full sequence alignment of *P. falciparum* (Pf3D7_0523800), *H. sapiens* (Uniprot P49281), and *S. cerevisiae* (Uniprot Q12078) DMT1 homologs. Asterisks denote fully conserved residues. Colons denote residues conserved between two proteins.

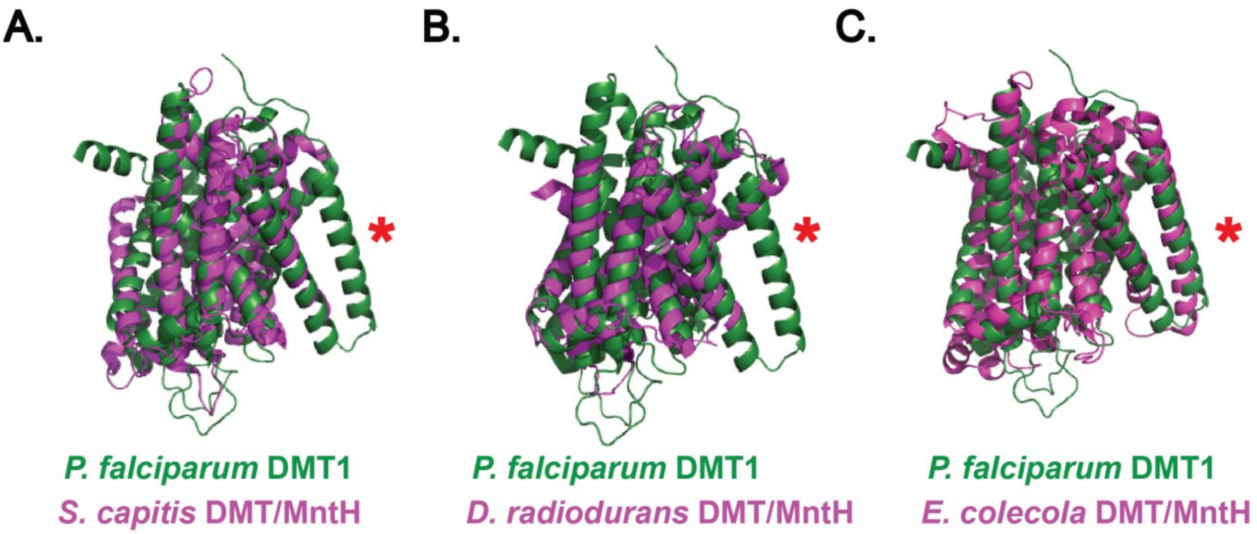


**Figure S2. Superposition of predicted AlphaFold structures of PfDMT1 with X-ray structures of homologs. (A)** *Staphylococcus capitis* (PDB 5M95), **(B)** *Deinococcus radiodurans* (PDB 5KTE), and **(C)** *Eremococcus coleocola* (PDB 5M87) crystal structures(35, 36). The disordered N-terminus of PfDMT1 was excluded from structural alignments. The red asterisk indicates the predicted $12^{th}$ transmembrane domain of PfDMT1. *S. capitis* DMT/MnTH and *D. radiodurans* DMT/MntH are 11 transmembrane domain proteins and *E. coli* DMT/MntH is a 12 transmembrane domain protein.

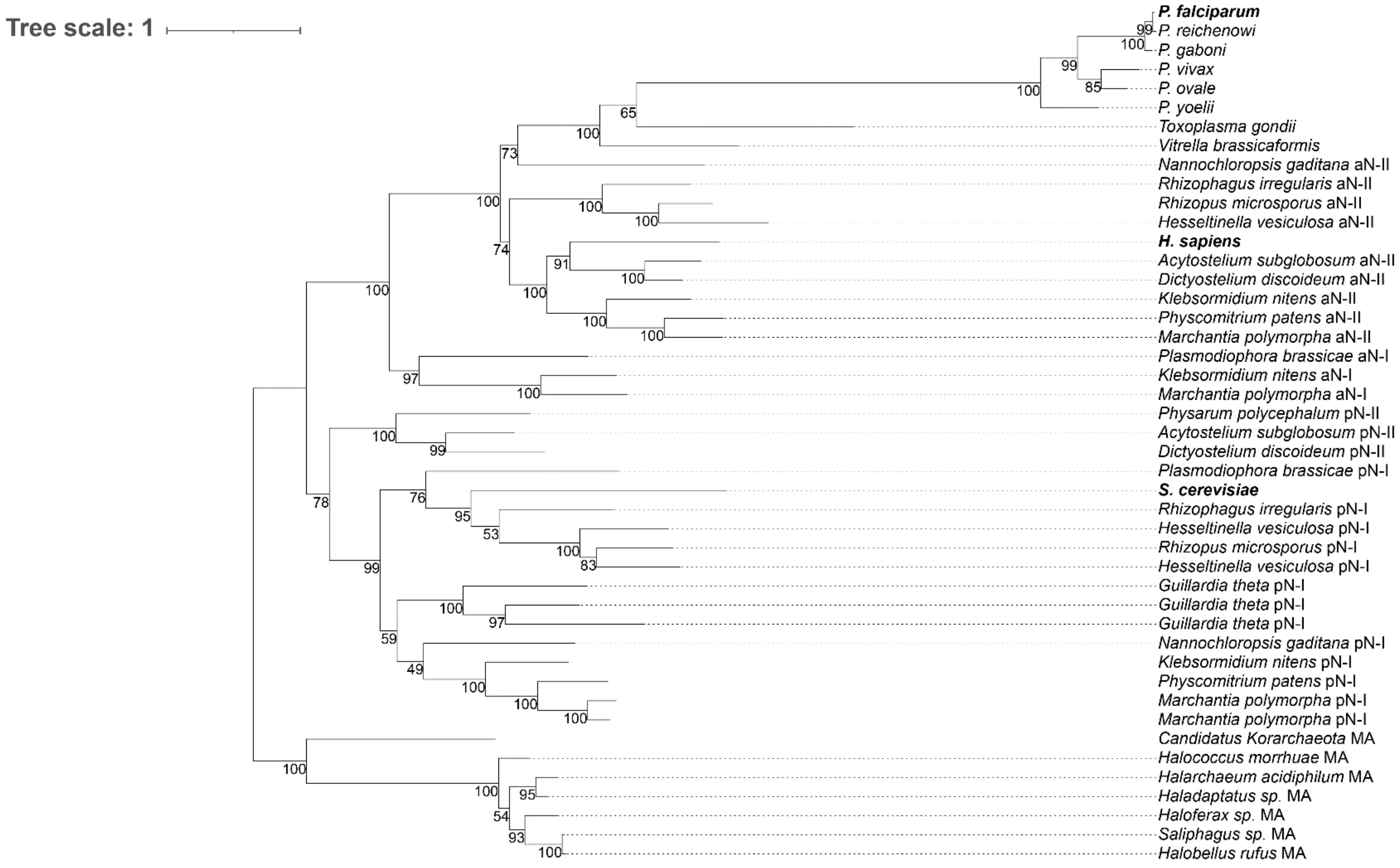


**Figure S3. Molecular phylogeny of PfDMT1.** All DMT1 subtype classifications (pN-I, pN-II, aN-I, aN-II) were previously assigned(40). Prokaryotic MntH (MA) transporters were used as the outgroup. Each node displays the calculated bootstrap values. The scale bar represents the expected sequence change per 100 bases.

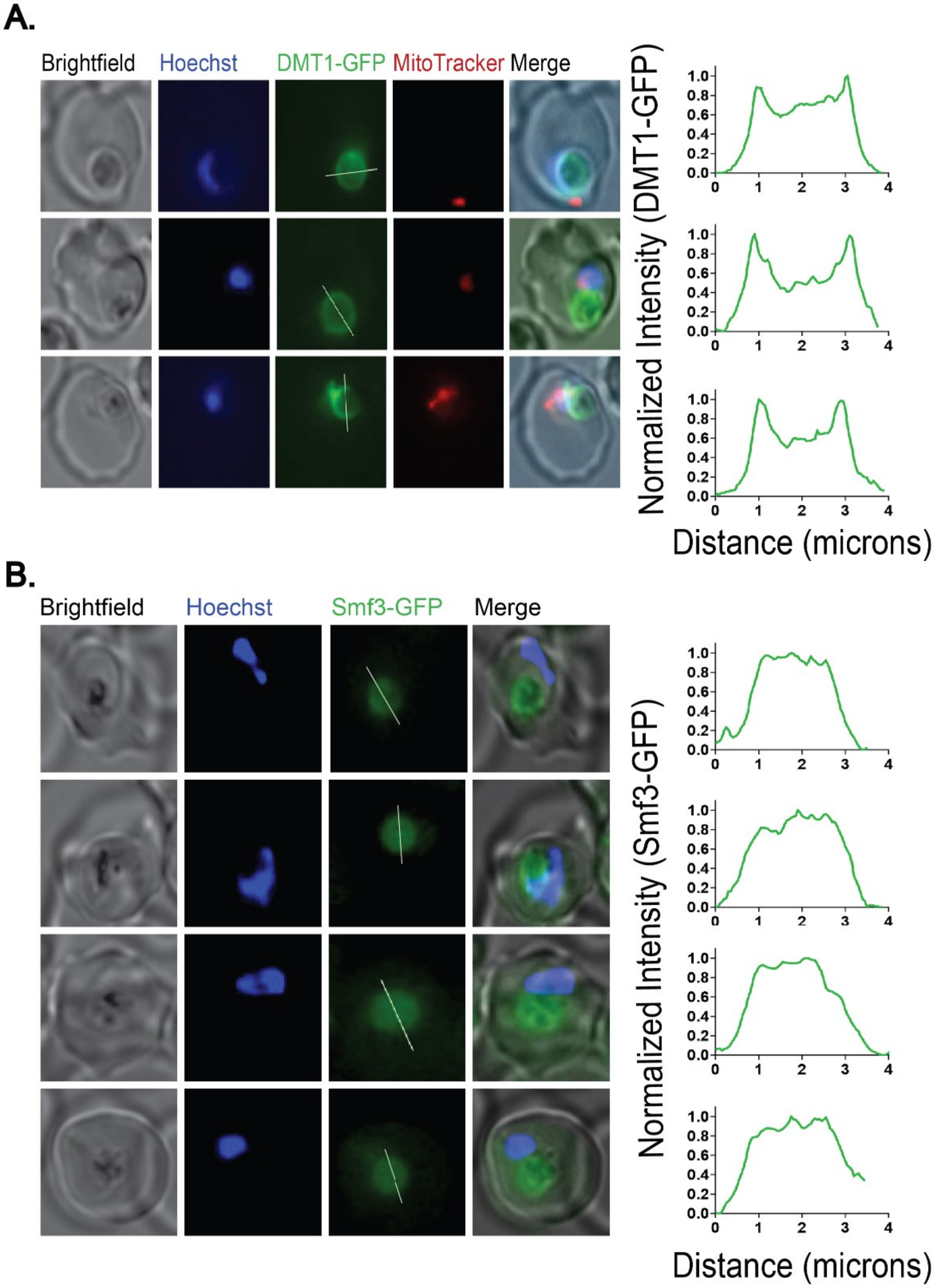


**Figure S4. Additional live-parasite microscopy images of DMT1 expression**. Live cell fluorescence microscopy of C-terminally tagged PfDMT1 **(A)** and Smf3 **(B)**. The intensity plots represent the normalized GFP fluorescence as a function of distance along the white line. Parasites in (A) were stained with 25 nM MitoTracker™ Red CMXRos.

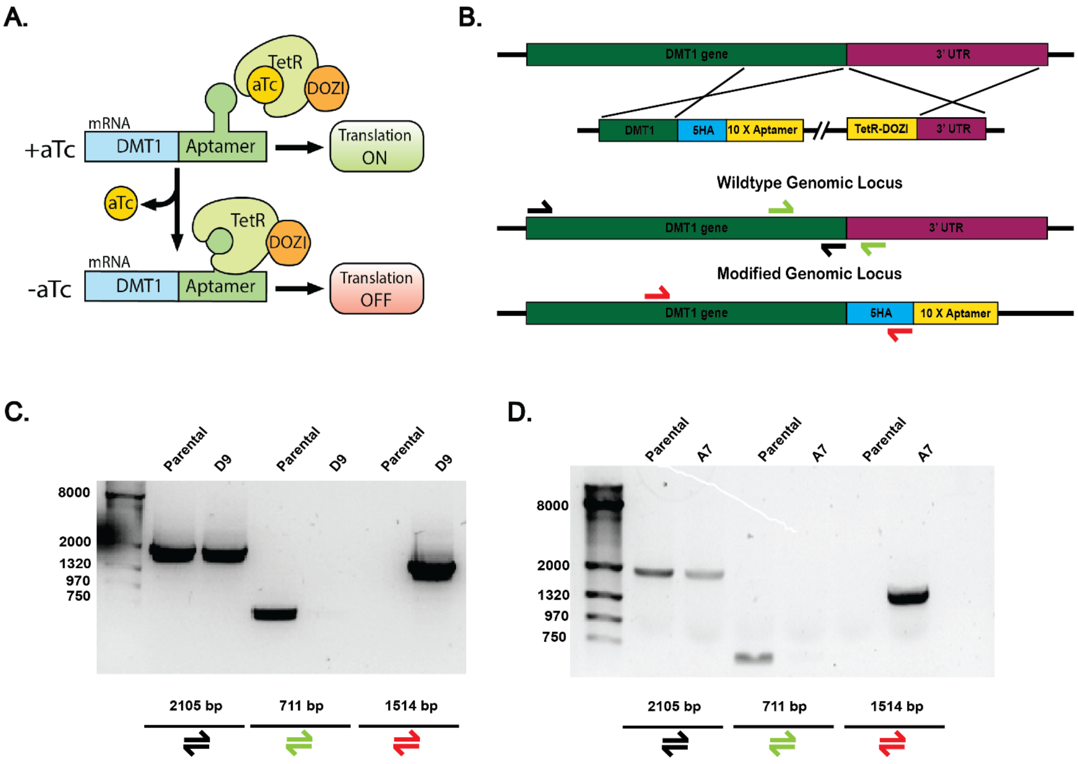


**Figure S5. Tagging PfDMT1 genomic locus for conditional knockdown**. **(A)** Schematic representation of aptamer/TetR-DOZI system for protein KD, reproduced from (cite Seyi's paper). **(B)** Scheme of strategy used to modify the endogenous PfDMT1 locus with the aptamer/TetR-DOZI system. **(C)** and **(D)** Agarose gels of PCR amplicons demonstrating successful modification of PfDMT1 locus in two clones from independent transfections of PfMev Nf54 *P. falciparum*. Color-coded primers indicate annealing sites with amplicon sizes.

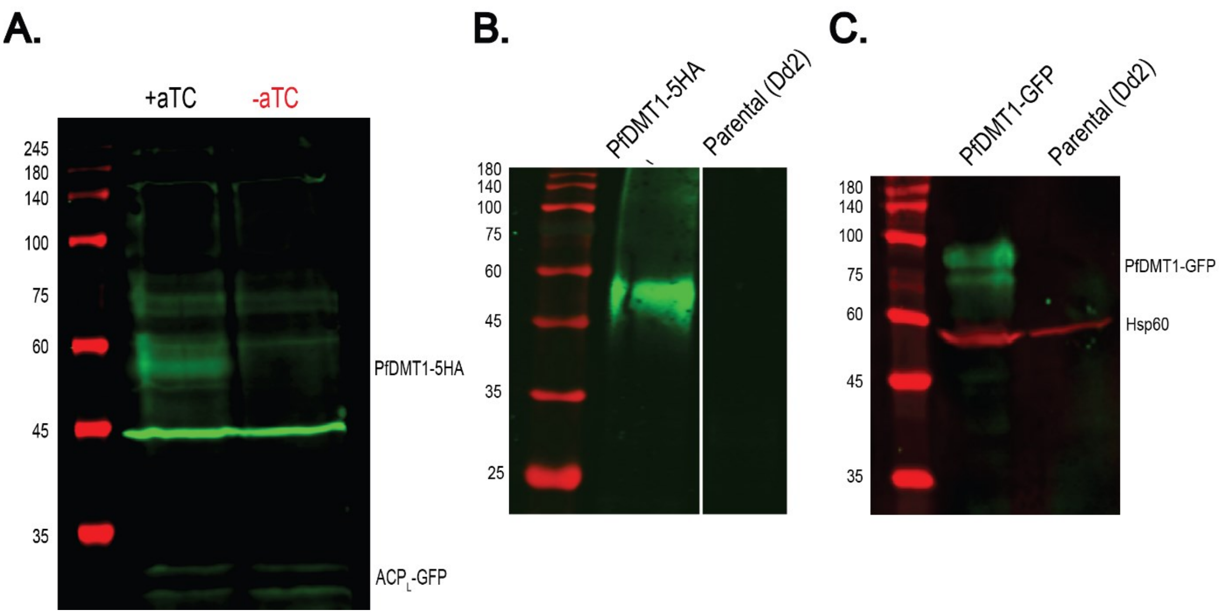


**Figure S6. Full SDS-PAGE/western blot analysis of PfDMT1 expression. (A)** Western blot analysis of endogenously tagged PfDMT1-5HA ± aTC. $ACP_L$-GFP present in the NF54 PfMev parasite line was used as a loading control(45). **(B)** Anti-HA immunoprecipitation and western plot analysis of PfDMT1-5HA and parental Dd2 parasites. **(C)** Western blot of episomally expressed PfDMT1-GFP and parental Dd2 parasites. Hsp60 was used as a loading control.

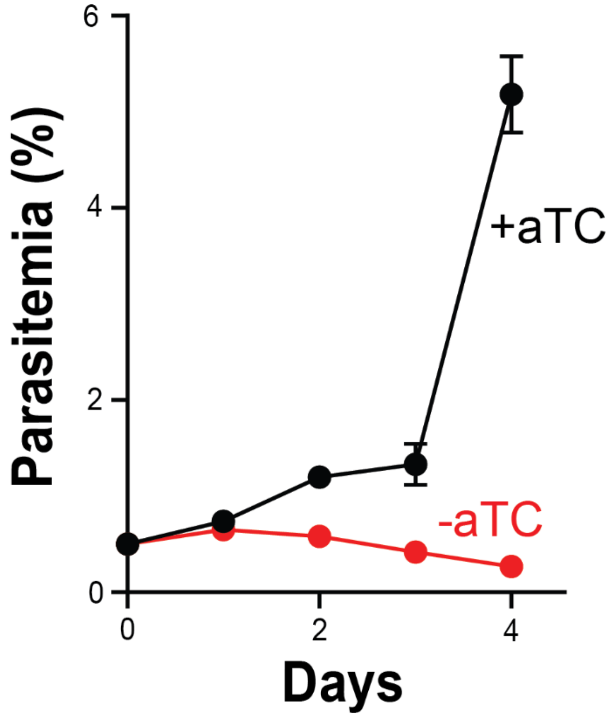


**Figure S7. Growth assay of PfDMT1 KD clonal line A7.** Data points and error bars are the average ± SD of biological triplicate measurements.

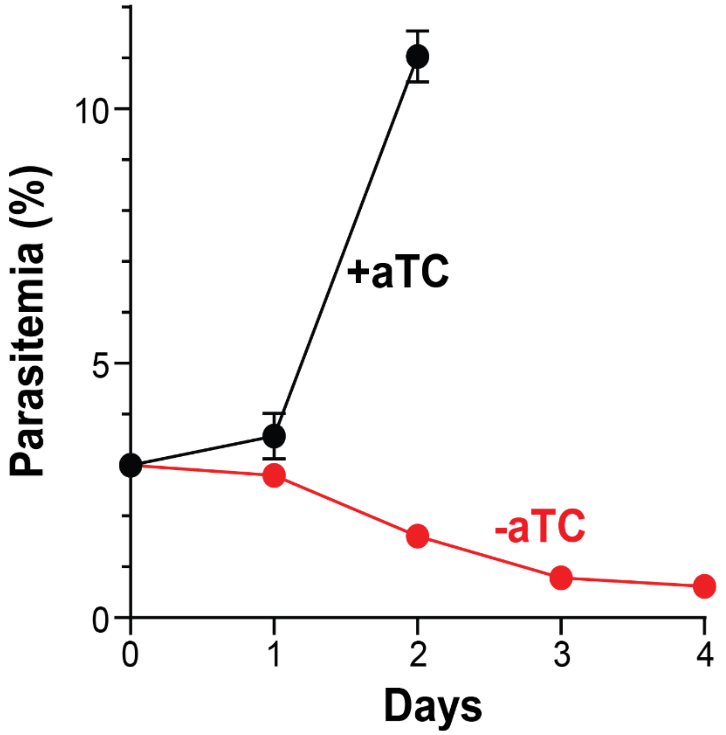


**Figure S8. Growth assay of PfDMT1 KD parasites (clone D9) seeded at a starting parasitemia of 3%.** Data points and error bars are the average ± SD of biological triplicate measurements.

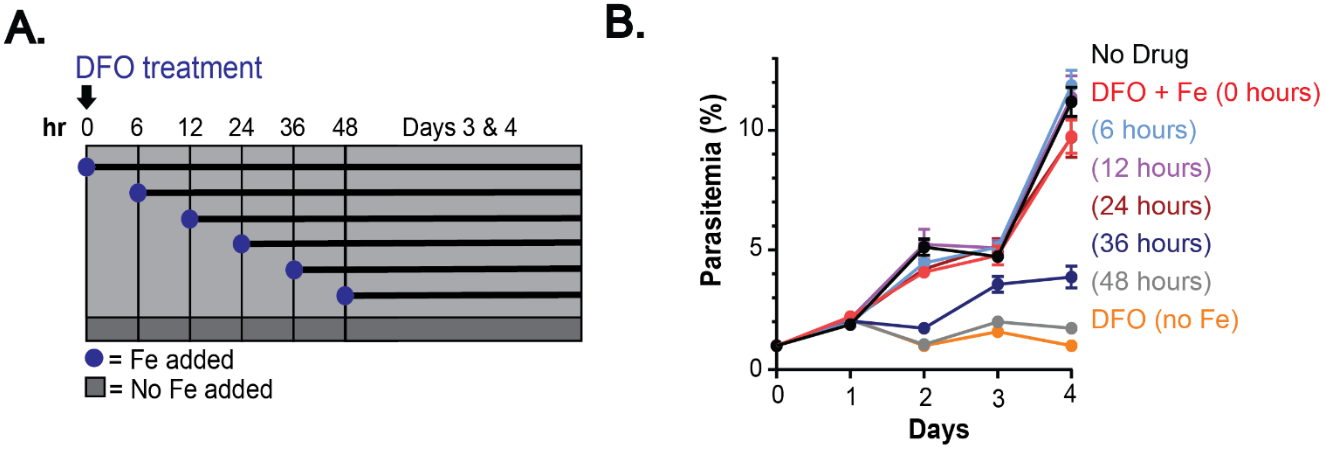


**Figure S9. Impacts of DFO treatment on parasite growth and rescue by exogenous iron. (A)** Schematic summary of delayed iron rescue of 60 μM DFO-treated parasites rescued with 60 μM $FeCl_2$. **(B)** Growth assay showing time dependence of $FeCl_2$ rescue after DFO treatment. Parasites were synchronized to a 4-hour window and treated with 60 μM of $FeCl_2$ at the indicated time points. Data points and error bars are the average ± SD of biological triplicate measurements.

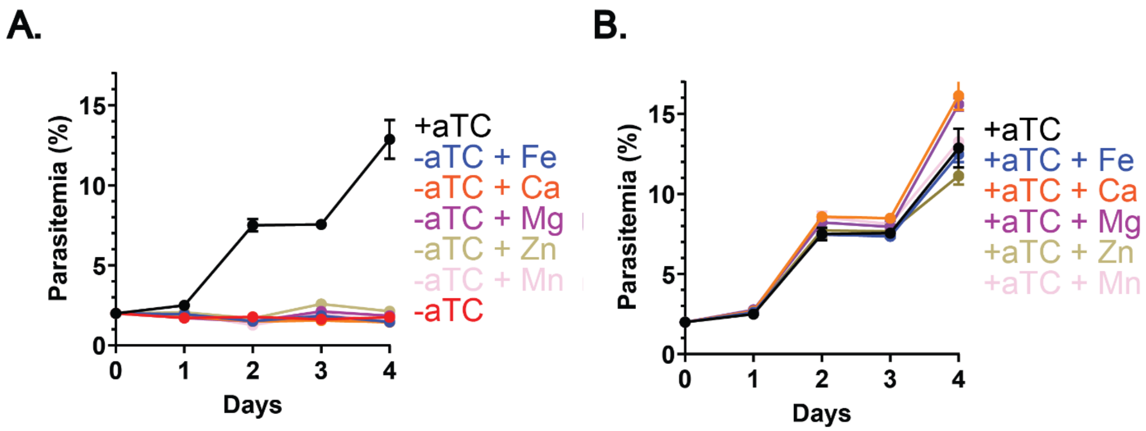


**Figure S10. Effect of exogenous metals on growth of PfDMT1 KD parasites. (A)** Growth assay of PfDMT1 KD parasites grown -aTC **(A)** +aTC **(B)** and treated with 125 μM $FeCl_2$, 125 μM $CaCl_2$, 125 μM $MgCl_2$, 5 μM $ZnCl_2$, or 20 nM $MnCl_2$. Data points and error bars are the average ± SD of biological triplicate measurements.

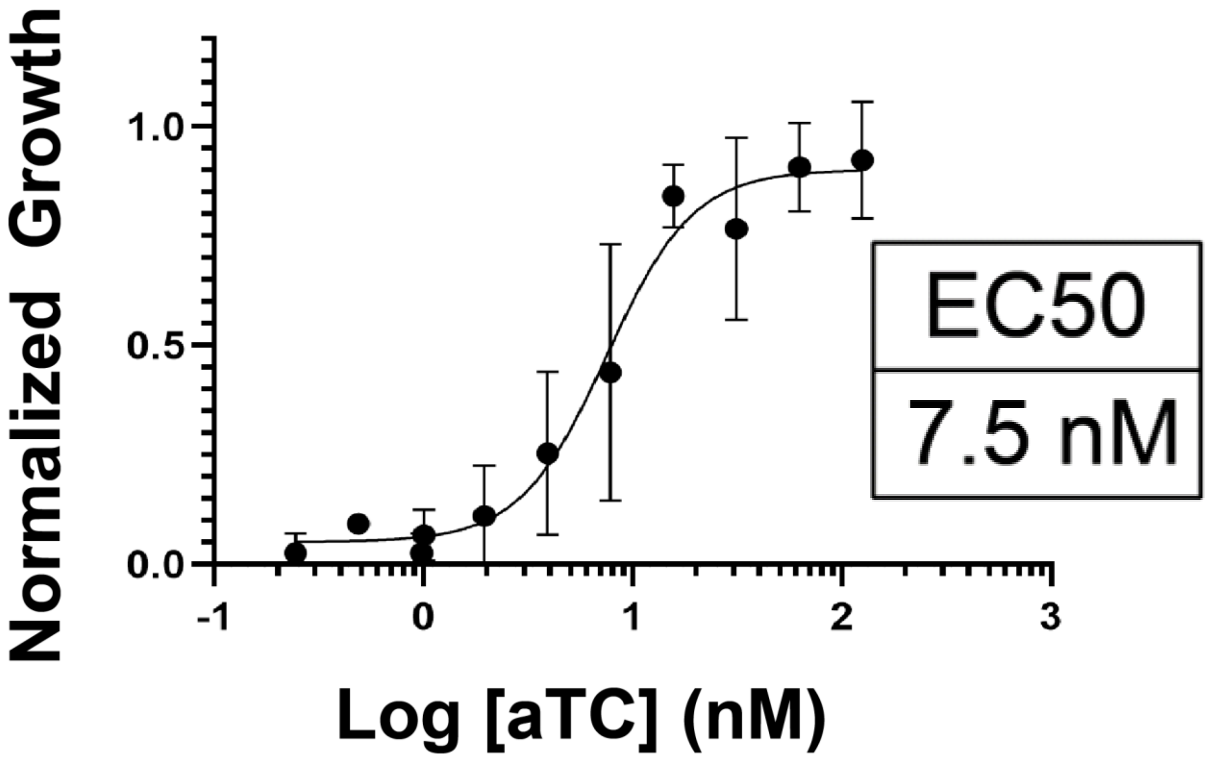


**Figure S11. Determination of aTC $EC_{50}$ value for PfDMT1 KD parasites**. Dose-response curve of parasite growth versus aTC concentration using PfDMT1 KD parasites. Log [aTC] nM indicates an $EC_{50}$ value = to 7.5 nM. Data points and error bars reflect the average ± SD of biological triplicates and were fit with a 3-parameter dose-response model in GraphPad Prism.

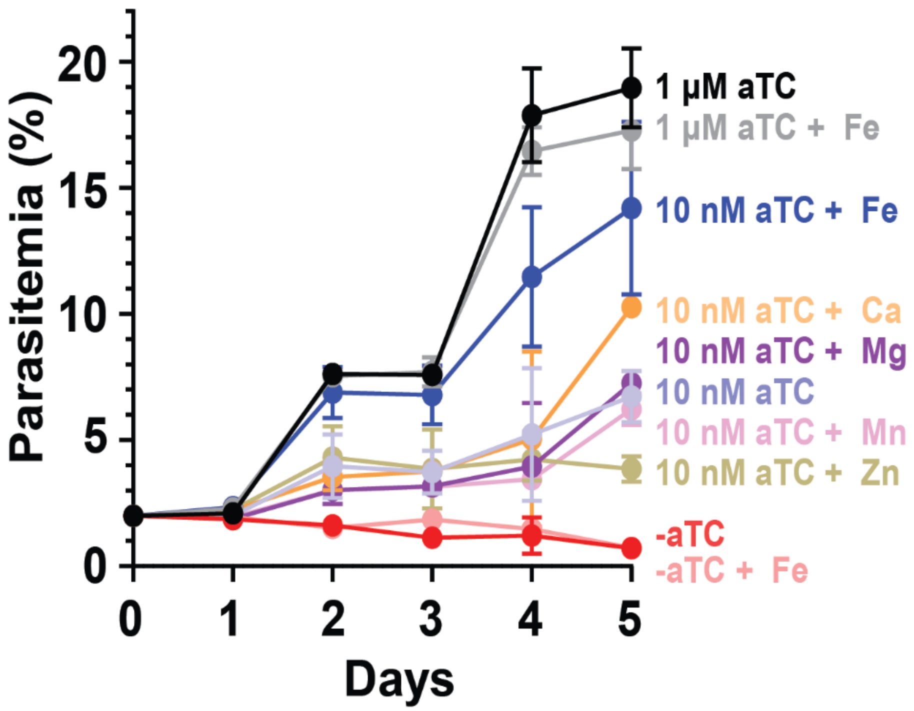


**Figure S12. Metal supplementation in intermediate PfDMT1 KD conditions.** Continuous growth assay of PfDMT1 KD parasites grown ±10 nM or 1 µM aTC ± 125 µM $FeCl_2$, $CaCl_2$, or $MgCl_2$, 20 nM $MnCl_2$ or 5 µM $ZnCl_2$. Growth assay data points represent the mean ± SD of 3-9 biological replicates.

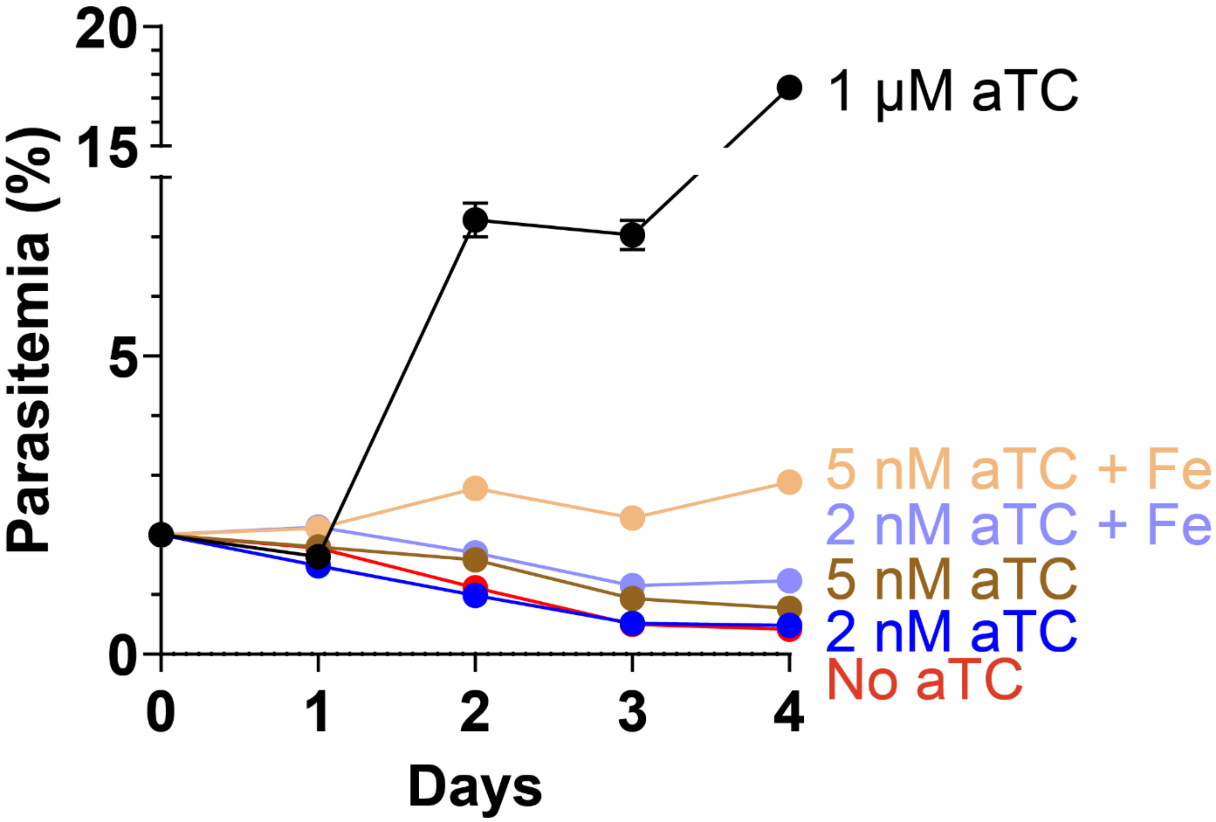


**Figure S13. Iron minimally rescues parasite growth at aTC concentrations below 10 nM.** Growth assay of PfDMT1 KD parasites grown at 5 nM or 2 nM aTC ± 125 µM $FeCl_2$. Data points and error bars are the average ± SD of biological triplicate measurements.

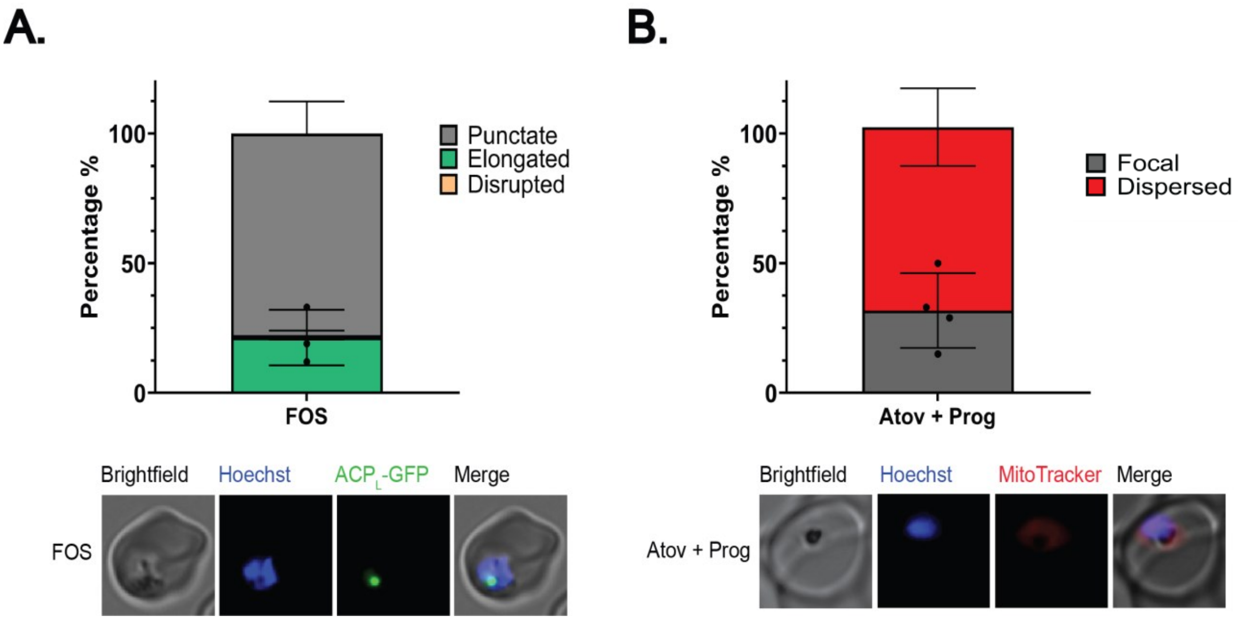


**Figure S14. Fosmidomycin and atovaquone/proguanil block apicoplast elongation and mitochondrial polarization. (A)** Top: Population analysis of apicoplast morphology in parasites treated with 10 µM fosmidomycin. Bottom: Live-cell images of fosmidomycin-treated PfDMT1 KD parasites. The apicoplast was visualized by $ACP_L$-GFP expressed in PfMev NF54 parasites. **(B)** Top: Population analysis of mitochondrial polarization in parasites treated with 100 nM atovaquone and 5 µM proguanil. Bottom: Live-cell images of PfDMT1 KD parasites stained with MitoTracker™ Red CMXRos after culture after treatment with atovaquone and proguanil. For (**A**) and (**B**) data is the mean of biological replicates where 12-20 parasites were imaged and scored. A two-tailed unpaired t-test was utilized to calculate the given p-values (full comparison given in Table S1 and S2). In all experiments, parasites were synchronized to a 4-hour window and treated at t=0 hours post-synchronization to rings.

| Experimental Conditions Compared: | p value: |
|---|---|
| + aTc vs. -aTC | 0.0001 |
| + aTc vs. DFO | 0.0003 |
| + aTc vs. Fos | <0.0001 |
| + aTc vs. WR | >0.9999 |
| -aTC vs. DFO | >0.9999 |
| -aTC vs. Fos | 0.4596 |
| -aTC vs. WR | 0.0001 |
| DFO vs. Fos | 0.0914 |
| DFO vs. WR | 0.0003 |
| Fos vs. WR | <0.0001 |

**Table S1.** Statistical comparison of all conditions assessed in Figure 4C. A two-tailed unpaired t-test was utilized to calculate the given p-values.

| Experimental Conditions Compared: | p value: |
|---|---|
| + aTc vs. + aTC + Proguanil | >0.9999 |
| + aTc vs. -aTC | >0.9999 |
| + aTc vs. -aTC + Proguanil | 0.0011 |
| + aTc vs. DFO | >0.9999 |
| + aTc vs. DFO + Proguanil | 0.0006 |
| + aTc vs. Atv + Proguanil | <0.0001 |
| + aTC + Proguanil vs. -aTC | >0.9999 |
| + aTC + Proguanil vs. -aTC + Proguanil | 0.003 |
| + aTC + Proguanil vs. DFO | >0.9999 |
| + aTC + Proguanil vs. DFO + Proguanil | 0.0017 |
| + aTC + Proguanil vs. Atv + Proguanil | <0.0001 |
| -aTC vs. -aTC + Proguanil | 0.0009 |
| -aTC vs. DFO | >0.9999 |
| -aTC vs. DFO + Proguanil | 0.0005 |
| -aTC vs. Atv + Proguanil | <0.0001 |
| -aTC + Proguanil vs. DFO | 0.0009 |
| -aTC + Proguanil vs. DFO + Proguanil | >0.9999 |
| -aTC + Proguanil vs. Atv + Proguanil | 0.3662 |
| DFO vs. DFO + Proguanil | 0.0005 |
| DFO vs. Atv + Proguanil | <0.0001 |
| DFO + Proguanil vs. Atv + Proguanil | 0.5987 |

**Table S2.** Statistical comparison of all conditions assessed in Figure 4E. A two-tailed unpaired t-test was utilized to calculate the given p-values.

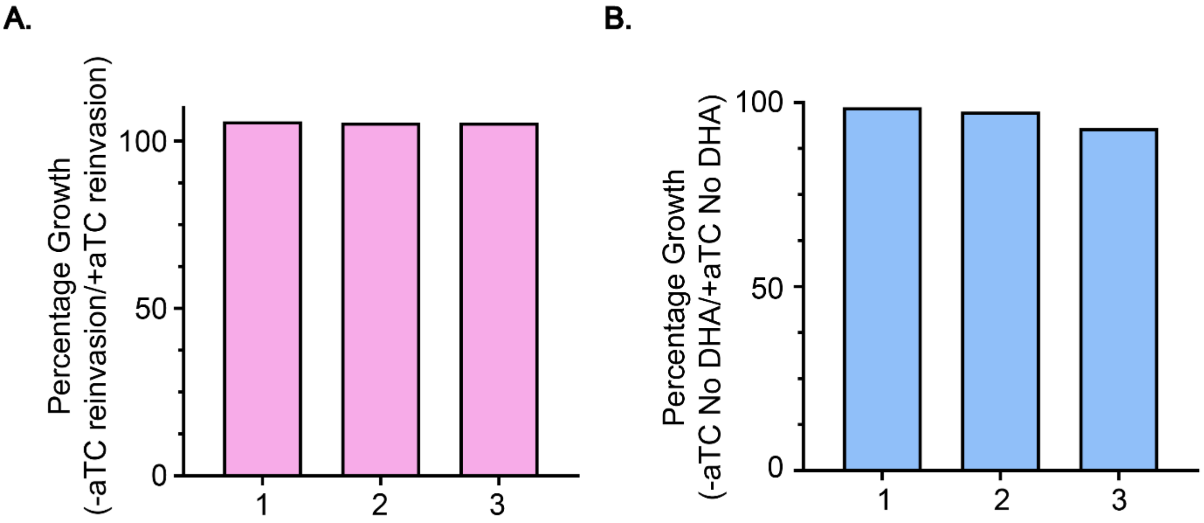


**Figure S15. Washout of aTC in schizonts does not impact their ability to rupture and invade new RBCs (A)** Comparison of parasite outgrowth when aTC was washed out of PfDMT1 parasites at 42 hours and allowed to reinvade. Each bar represents one biological replicate normalized to growth in +aTC conditions. **(B)** Relative parasite growth of PfDMT1 KD parasites that were not pulsed with 700 nM DHA and grown for 6 hours -aTC versus +aTC in the ring-stage survival assay. After 6 hours all parasites were grown in +aTC media for 66 hours until parasitemia was measured. Each bar represents one biological replicate.

A.

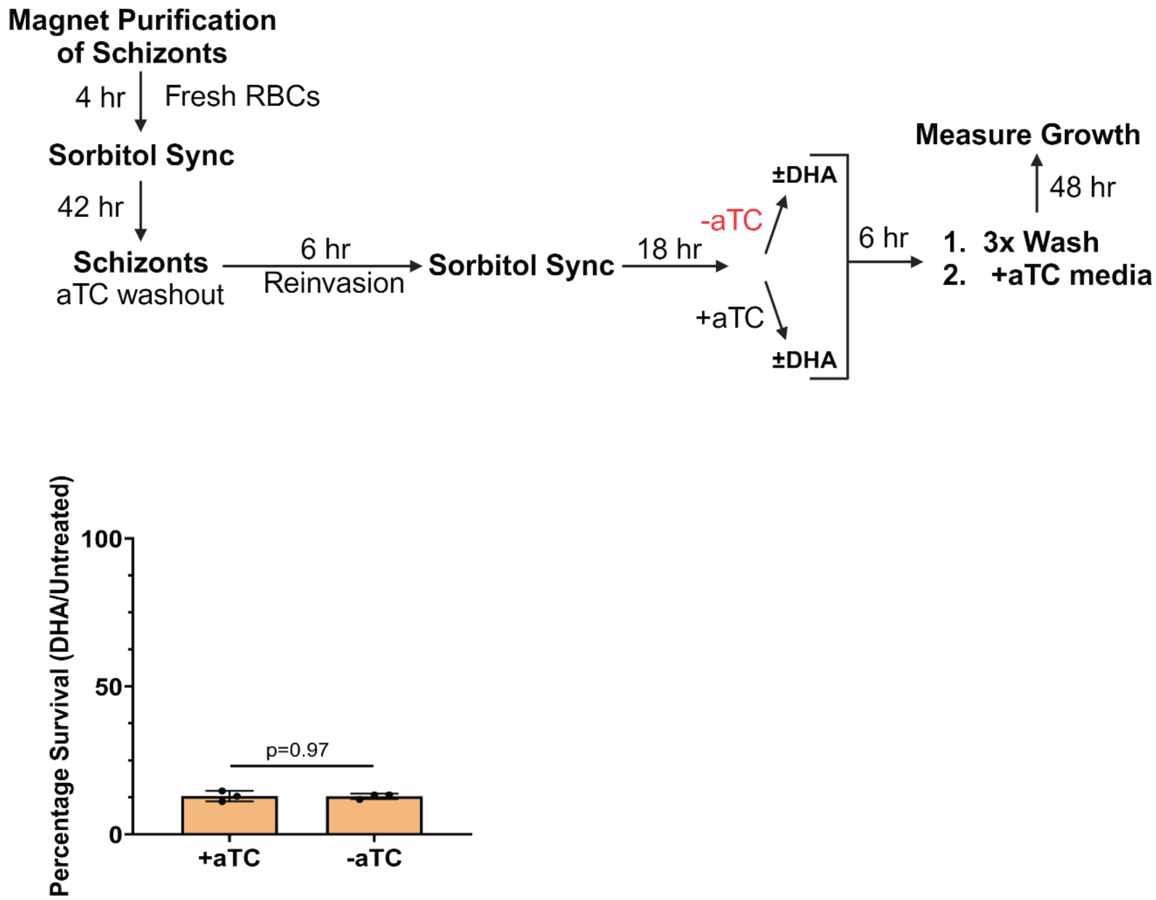


**Figure S16. Loss of PfDMT1 does not alter parasite sensitivity to artemisinin in trophozoite-stage parasites. (A)** Schematic summary of assay modified from the RSA in Fig. 5A. Trophozoite PfDMT1 KD parasites were growth ±aTC for 18 hours and pulsed with 700 nM DHA before drug washout and monitoring of growth in +aTC conditions. **(B)** Survival of trophozoite PfDMT1 KD parasites pulsed with 700 nM DHA for 6 hours. Each data point represents the growth percentage of DHA-treated versus untreated parasites outgrown for 48 hours after DHA exposure. Percentages are the mean ± SD of biological triplicate assays analyzed by two-tailed unpaired t-test.
